# Rab7a is required to degrade select blood-brain barrier junctional proteins after ischemic stroke

**DOI:** 10.1101/2023.08.29.555373

**Authors:** Azzurra Cottarelli, Danny Jamoul, Mary Claire Tuohy, Sanjid Shahriar, Michael Glendinning, Grace Prochilo, Aimee L. Edinger, Ahmet Arac, Dritan Agalliu

## Abstract

Adherens (AJ) and tight junction (TJ) integrity is critical for blood-brain barrier (BBB) function in the healthy brain. Junction disassembly due to degradation of AJ and TJ proteins leads to acute BBB dysfunction after ischemic stroke, but the mechanisms are not fully understood. Here, we show that endothelial cell deletion of Rab7a, a small GTPase crucial for protein degradation through the endolysosomal system, reduces acute BBB dysfunction and improves neuronal health in mice after ischemic stroke by preventing degradation of select junctional proteins and preserving TJ structural morphology. Two pro-inflammatory cytokines, TNFα and IL1β, that trigger barrier disruption in brain endothelial cells (BECs) *in vitro* and are upregulated in stroke, contribute to Rab7a activation. Silencing Rab7a *in vitro* partially rescues cytokine-driven barrier disruption in BECs by reducing internalization of some junctional proteins and the formation of F-actin bundles at cell junctions. Rab7a is, therefore, critical for degradation of select junctional proteins during the acute BBB damage after ischemic stroke.

## INTRODUCTION

The stability of adherens (AJs) and tight junction (TJs) strands formed between brain endothelial cells (BECs) lining the blood vessels is critical for blood-brain barrier (BBB) integrity and together with low rates of transcellular transport, restricts the permeability of blood-derived proteins and immune cells into the healthy central nervous system (CNS; reviewed in ^1^). Disassembly of TJ strands associated with BBB damage occurs in several neurological disorders including ischemic stroke, a complex and devastating neurological condition that is the second leading cause of death and the third leading cause of disability worldwide (reviewed in ^2^). In addition to the acute damage triggered by BBB breakdown after ischemic stroke (reviewed in ^3^), BECs that survive and proliferate in the peri-infarct region of the brain, but do not maintain BBB properties, facilitate further tissue damage and exacerbate the clinical prognosis of stroke leading to post-stroke cognitive dementia ^4–6^. Elucidating the mechanisms that promote stabilization of BBB junctions is critical to identify targets that could improve both short- and long-term stroke outcomes. Yet, how AJ and TJ proteins are degraded after ischemic stroke to promote acute BBB damage is not fully understood.

During the reperfusion phase of acute ischemic stroke, there is a biphasic increase in BBB permeability that contributes to vasogenic edema, hemorrhage and increased mortality ^7–10^. Although BEC junctional strands are stable in the healthy brain ^11, 12^, we have shown using green fluorescent protein (GFP)-tagged Claudin-5 that BBB TJs become highly dynamic and exhibit structural abnormalities between 24-58 hours after transient middle cerebral artery occlusion (t-MCAO), a rodent model for ischemic stroke ^11^. This phase coincides with increased expression of several pro-inflammatory cytokines and recruitment of immune cells into the CNS (reviewed in ^13, 14^), that contribute to stroke pathophysiology, including further BBB damage, and worse clinical outcomes ^15–17^. What mechanisms drive BBB AJ and TJ disassembly after ischemic stroke? Increased expression and activity of several matrix metalloproteases (MMPs; e.g., MMP2 and MMP9) after ischemic stroke contributes to acute BBB damage by cleaving the extracellular domains of several adherens (VE-Cadherin) and tight (Claudin-5, Occludin) junction proteins in BECs (reviewed in ^18, 19^). Ubiquitin-mediated proteasome degradation has also been proposed as a mechanisms for degradation of transmembrane proteins (VE-Cadherin, Claudin-5 and Occludin) and intracellular adaptors (ZO-1 and β-catenin) that anchor transmembrane junctional proteins to the cytoskeleton in various diseases including ischemic stroke (reviewed in ^20, 21^). Finally, caveolae have also been proposed to promote internalization and degradation of TJ transmembrane proteins such as Claudin-5 and Occludin in brain and peripheral ECs ^22^. However, *Caveolin-1 (Cav-1)*^−/−^ mice lacking caveolae ^23, 24^ show a similar degree of paracellular BBB damage and similar infarct size to wild-type mice after t-MCAO ^11, 25^, suggesting a caveolar-independent mechanism for BBB TJ degradation.

Enhanced endocytotic degradation of transmembrane and intracellular BBB junctional proteins in response to the inflammatory milieu is another potential mechanism for persistent AJ and TJ strand disassembly post-ischemic stroke. EC junctional strand protrusions express the early endosomal marker EEA-1 ^12^ and they are inhibited by treatment with inhibitors of endocytosis ^26^. During endocytosis, early endosomes and their cargoes can either be sorted back to the plasma membrane as recycling endosomes, or directed to degradation by maturing into late endosomes that ultimately fuse with lysosomes ^27^. The small GTPase Rab7a is critical for sorting cargoes into the late endosome, biogenesis of lysosomes and phagocytosis, as well as regulation of substrate degradation, antigen presentation, cell signaling/survival and microbial infection ^28, 29^. Although the endolysosomal functions of Rab7 have been studied in healthy mammalian cells, how its activity is regulated in neurological diseases is not well understood. Growth factor deprivation in cancer cells increases the fraction of Rab7a associated with membranes, and Rab7a-GTP (active form) triggers cell death due to degradation of nutrient transporters essential for survival ^30–32^. Conversely, Rab7a inhibition after growth factor withdrawal promotes cancer cell survival due to recycling of transporters to the cell surface ^33, 34^. Regulatory proteins that control Rab7a activity have been difficult to identify in many cells. TBC1D15 has been identified as a putative Rab7a inactivator (GAP), since it accelerates GTP hydrolysis by Rab7a *in vitro* and its overexpression disrupts lysosomal morphology and blocks growth factor withdrawal-induced death in cancer cells ^33^. Ccz1, a guanidine exchange factor (GEF) that acts together with vacuolar fusion protein Mon1, has been shown to promote Rab7a activation and Rab7a-mediated membrane fusion in yeast ^35^ and eukaryotic cells ^36–39^. But, it is unclear whether Mon1/Ccz1 complex activates Rab7a in ECs and the role of Rab7a in EC function has not been fully elucidated. Rab5a and Rab7a regulate VEGFR2 receptor trafficking and the response of ECs to VEGF-A ^40^ via the Rab effector protein RABEP2 ^41^. The liver kinase B1 (LBK1) has also been identified as a Rab7a effector in ECs that promotes Neuropilin-1 receptor trafficking and degradation to inhibit angiogenesis ^42^. Moreover, Rab4 activation and Rab9 inhibition are also critical for the vascular permeability in lung ECs, through regulation of VE-Cadherin localization to cell junctions ^43^. Yet, how endothelial Rab7a contributes to acute BBB dysfunction after ischemic stroke is unknown.

Here, we show that endothelial-specific deletion of Rab7a improves neuronal survival and reduces acute BBB disruption in mice after ischemic stroke by preventing degradation of some junctional proteins and preserving TJ morphology at the electron microscopy level. TNFα and IL1β, two pro-inflammatory cytokines that are upregulated in the brain after ischemic stroke ^14^ induce Rab7a activation in BECs *in vitro*. Silencing Rab7a *in vitro* partially rescues cytokine-driven barrier disruption by reducing internalization of some junctional proteins and formation of F-actin bundles in response to TJ structural disassembly at cell borders. These findings identify Rab7a as a critical regulator of select junctional protein degradation during the acute BBB damage following ischemic stroke.

## RESULTS

### Rab7a deletion in endothelial cells reduces acute paracellular BBB permeability 48 hours after t-MCAO

Rab7a inhibition is critical to prevent degradation of nutrient transporters and support survival in lymphocytes and cancer cells ^31, 33, 34^. To determine if Rab7a may similarly mediate degradation of BBB TJ proteins in BECs after ischemic stroke, we generated a Rab7a-inducible EC knockout mouse strain [*Rab7a^iECKO^ (Rab7a^fl/fl^; VEC-Cre^ERT2+/−^; eGFP::Claudin5^+/−^*] to ablate Rab7a in ECs upon administration of 4-OH-tamoxifen (**Figure 1A**). These mice were crossed to eGFP::Claudin5 transgenic mice to visualize BBB TJ strand morphology after ischemic stroke ^11^. We confirmed the efficiency and specificity of Rab7a elimination in BECs by decreased Rab7a protein expression in Lectin (BSL)^+^ cortical blood vessels, but not other CNS cell types, of *Rab7a^iECKO^* compared to *Rab7a^fl/fl^* (*Rab7a^fl/fl^; eGFP-Claudin5^+/−^*; referred as WT) cortices by immunofluorescence (**Figures 1B-C”**). We also measured the efficiency of 4-OH tamoxifen-induced Cre deletion in BECs by crossing the *VEC-Cre^ERT2+/−^* strain with an Ai14 reporter mouse strain and found that all CNS blood vessels expressed tdTomato after 4-OH tamoxifen administration (**Figure 1D-F”**, yellow arrows).

**Figure 1.**
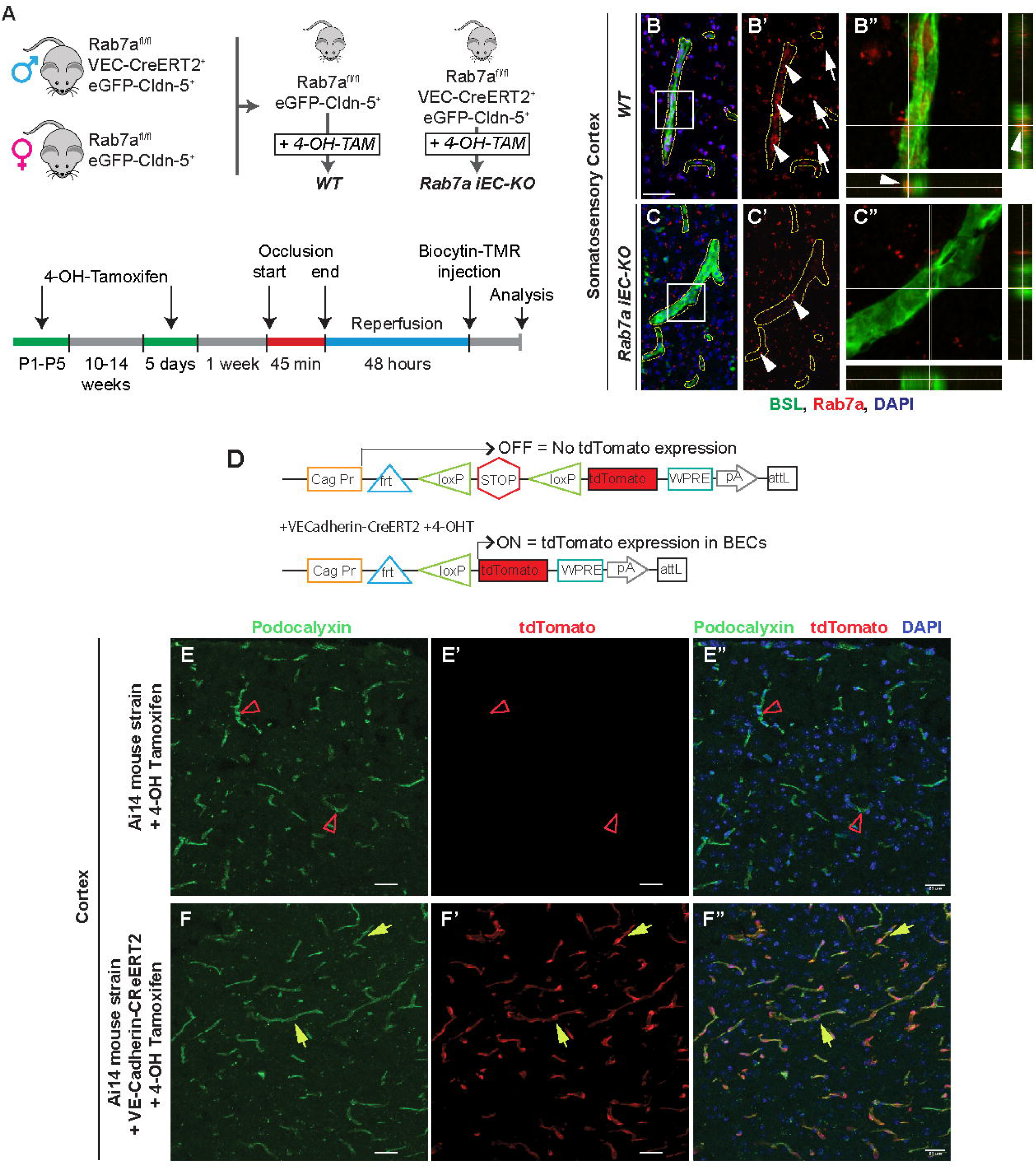
Generation of endothelial-specific Rab7a deficient mice. (**A**) Diagrams of the breeding strategy to generate the inducible endothelial Rab7a knockout (*Rab7a^iECKO^*) mice and the experimental setup to analyze blood-brain barrier (BBB) permeability after t-MCAO. (**B-C”**) Immunofluorescence images for Griffonia (Bandeiraea) Simplicifolia Lectin I (BSL, green), Rab7a (red) and DAPI (blue) in the cortex of healthy *Rab7a ^fl/fl^* (WT, **B-B”**) and *Rab7a^iECKO^* (**C-C”**) mice. Cortical vessels are delineated by yellow dotted lines (**B’**, **C’**). Boxed areas are magnified in the orthogonal view images on the right (**B”**, **C”**). Arrowheads indicate endothelial Rab7a, arrows indicate non-endothelial Rab7a. There is no residual Rab7a protein in brain endothelial cells of mutant mice. Scale bar = 30 μm. **(D)** Schematic diagram of Ai14 ROSA reporter mice containing a *loxP-*flanked STOP site before tdTomato to visualize the Cre-mediated recombination efficiency following breeding with the *VE-Cadherin-Cre^ERT2^* mice and 4-OH tamoxifen administration. **(E-F’)** Immunofluorescence images for Podocalyxin (vessel marker, green), tdTomato (red), and DAPI (blue) in the cortex of Ai14 mice and Ai14 + *VE-Cadherin-Cre^ERT2^*mice show that tdTomato is expressed in all blood vessels following Cre-mediated recombination induced by 4-OH tamoxifen. Normal green arrowhead corresponds to vessels expressing tdTomato, red open arrowhead indicates vessels not expressing tdTomato. Scale bar = 23 µm.

We induced ischemic stroke in *Rab7a^fl/fl^* and *Rab7a^iECKO^* mice by performing t-MCAO for 45 minutes ^11^ and analyzed the effect of Rab7a EC-specific elimination on acute changes in paracellular BBB permeability at 48 hours after ischemic stroke following injection of a small dye biocytin-TMR (890 Da) dye that preferentially extravasates from blood vessels through the paracellular route^11, 12^. *Rab7a^fl/fl^* mice [referred to as wild-type (WT) mice] showed an intense biocytin-TMR fluorescence signal in the ipsilateral cortex and striatum in all seven bregma regions after stroke (**Figure 2A, B**). In contrast, the extravascular tracer leakage was less frequent and extensive in the ipsilateral cortex and striatum of matched bregma regions from *Rab7a^iECKO^* mice (**Figure 2C, D**). The average leakage area, reflective of BBB disruption, was significantly reduced in *Rab7a^iECKO^* compared to WT brains across all bregmas, corresponding to ~2.5 fold decrease in the overall volume of tracer leakage (**Figure 2E, F**). In addition, the fraction of the brain area affected by BBB impairment was reduced ~2.0-2.5 fold in some bregma regions of *Rab7a^iECKO^* mice, and the amount of tracer leakage, measured as mean fluorescence intensity of the extravasated dye in the parenchyma, was also significantly reduced in *Rab7a^iECKO^* mice (**Figure 2F, G**). Thus, Rab7a EC-specific deletion rescues the acute increase in paracellular BBB permeability at 48 hours after t-MCAO.

**Figure 2.**
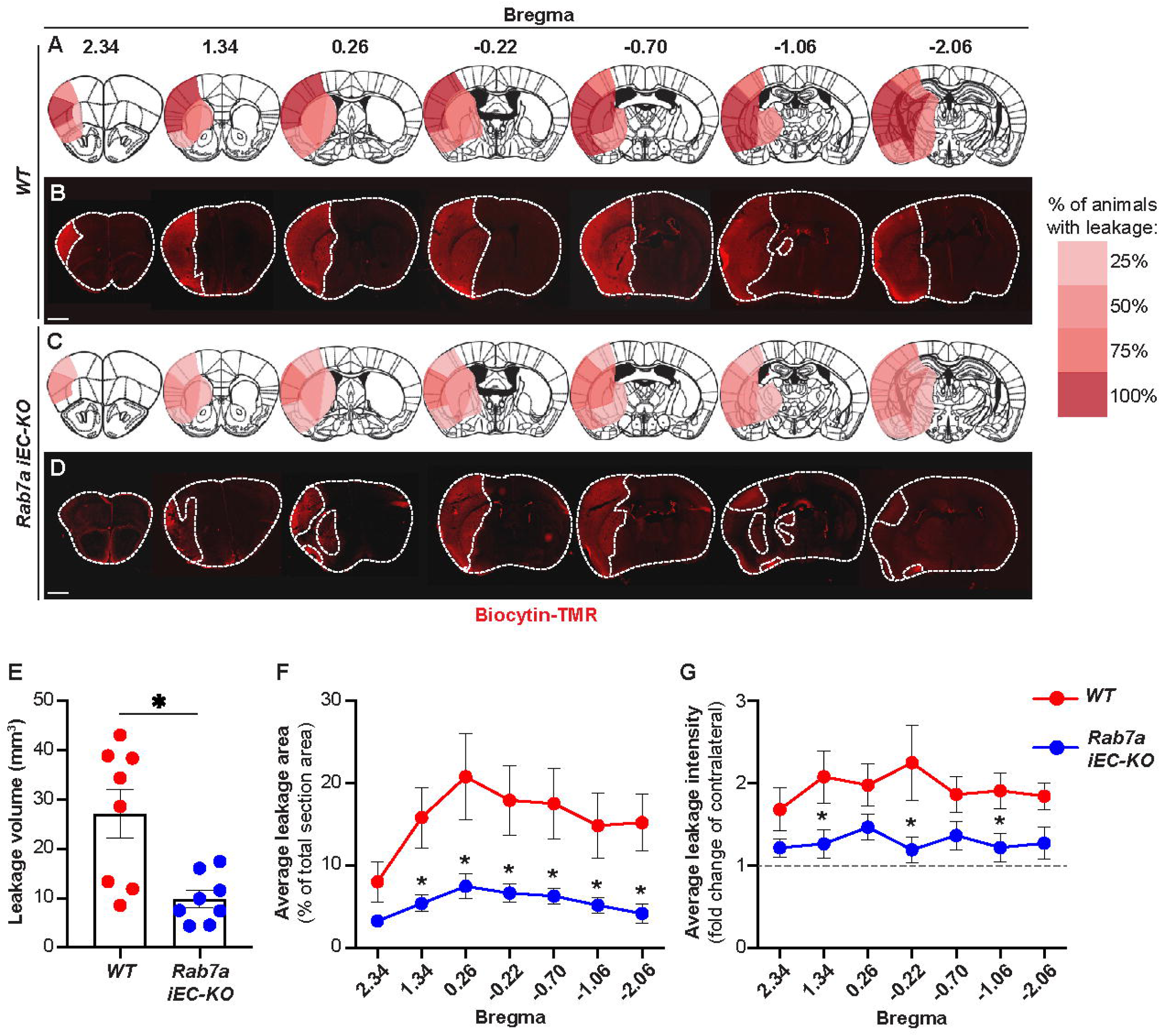
*Rab7a^iECKO^* mice show a partial rescue in the paracellular BBB permeability 48 hours after t-MCAO. (**A-D**) Fluorescent micrographs and heatmaps showing biocytin-TMR tracer extravasation in seven distinct brain regions in WT (**A, B**) and *Rab7a^iECKO^* (**C, D**) mice 48 hours after t-MCAO (biocytin-TMR was injected 30-45 minutes before analysis). Dotted lines outline the border of the brain section and the leakage area. The heatmaps show the fraction of animals displaying BBB leakage in each region represented as a scale (0-100%) of red hues. (**E**) Quantification of biocytin-TMR leakage volume in the brain of WT and *Rab7a^iECKO^* mice 48 hours after t-MCAO (n=8 animals/group). Data are means ± s.e.m. *: p<0.05; Student’s t-student. (**F**) Quantification of biocytin-TMR leakage area in each bregma region from WT and *Rab7a^iECKO^* mice 48 hours after t-MCAO. Each dot represents the average of n=8 animals / group. Data are means ± s.e.m. *: p<0.05; one-way ANOVA with post-hoc Tukey’s correction. (**G**) Quantification of theintensity of biocytin-TMR leakage in the ipsilateral cortex of each bregma region from WT and *Rab7a^iECKO^* mice 48 hours after t-MCAO. The dotted line represents the average fluorescence intensity in the contralateral cortex. Each dot represents the average of n=8 animals. Data are means ± s.e.m. *: p<0.05; one-way ANOVA with post-hoc Tukey’s correction.

To determine if Rab7a deletion in ECs impacts similarly changes in transcellular BBB permeability after ischemic stroke, we analyzed the leakage of endogenous serum immunoglobulin G (IgG) and a 70kDa dextran-TMR administered intravenously at 48 hours after t-MCAO. No significant differences in either the volume, area, or fluorescence intensity of serum IgG leakage were observed between the WT and *Rab7a^iECKO^* brains (**Figure 3A-G**). However, we could not detect any 70 kDa dextran-TMR leakage in the ipsilateral cortex of both genotypes at 48 hours after t-MCAO after extraction of the tracer from the brain and measurement of fluorescence intensity (**Figure 3H**), suggesting that the BBB was not leaky to 70 kDa dextran-TMR at 48 hours after t-MCAO. It is likely that the serum IgG leakage reflects BBB leakage at an earlier timepoint after t-MCAO. Correspondingly, the levels of Caveolin-1 (Cav-1) protein that mediates transcellular transport across ECs, were similarly increased in total brain lysates from the ipsilateral cortex of *Rab7a^fl/fl^* (WT) and *Rab7a^iECKO^* mice 48 hours after t-MCAO (**Figure 3I, J**), confirming that Rab7a EC-specific deletion did not affect the acute transcellular BBB leakage after ischemic stroke. Thus, Rab7a EC-specific deletion rescues significantly the acute changes in paracellular, but not transcellular, BBB permeability after ischemic stroke.

**Figure 3.**
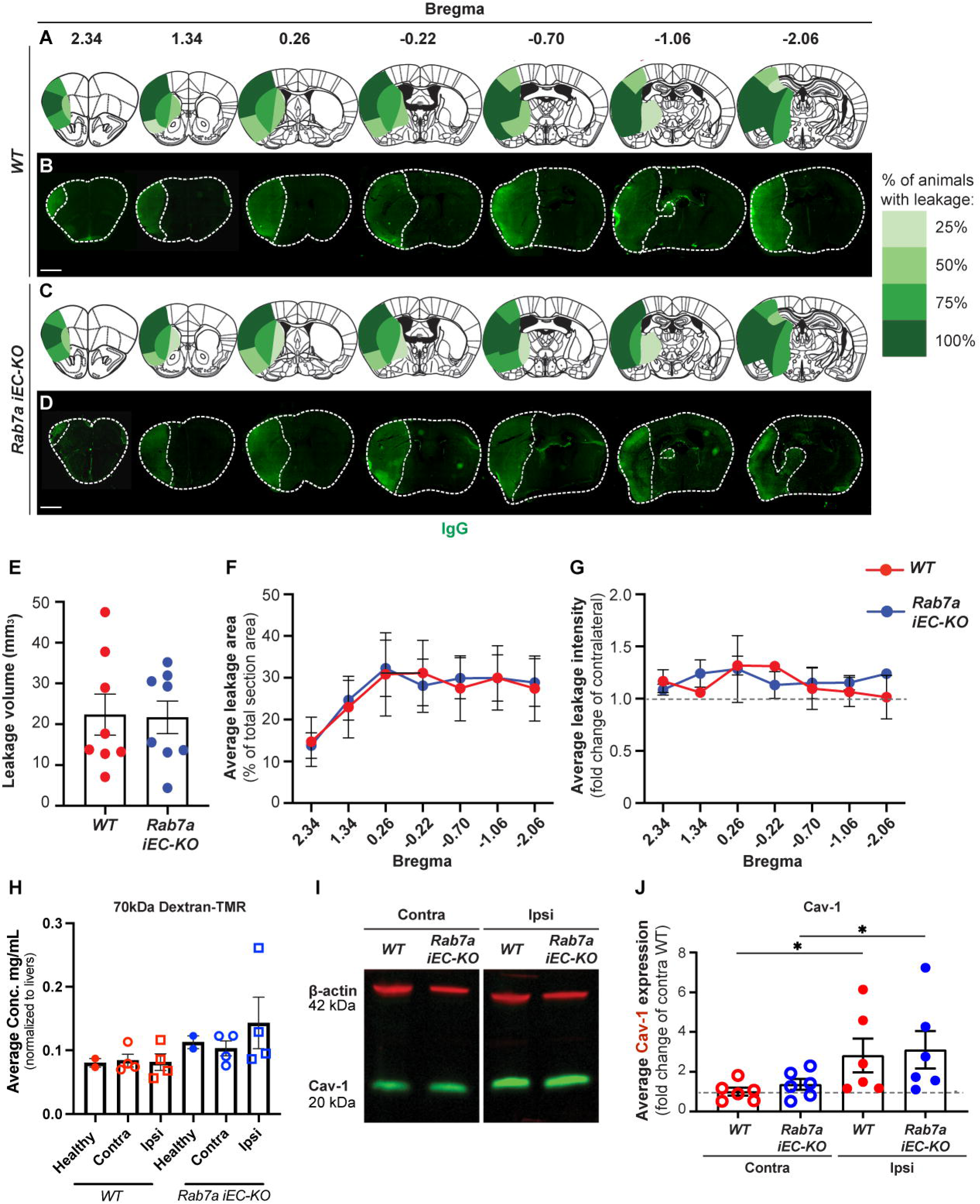
Rab7a elimination in brain endothelial cells has no effect on the increase in transcellular BBB permeability 48 hours after t-MCAO. (**A-D**) Fluorescent micrographs and heatmaps showing serum IgG extravasation into seven distinct brain regions of WT and *Rab7a^iECKO^* mice 48 hours after t-MCAO. The brain sections were stained for mouse Immunoglobulin G (IgG; green) to detect leakage of serum IgG from the blood vessels into the brain parenchyma (dotted lines outline the borders of the brain section and serum IgG leakage area). Heatmaps show the fraction of animals displaying serum IgG leakage in each brain region represented as a scale (0-100%) of green hues. (**E**) Quantification of serum IgG leakage volume in WT and *Rab7a^iECKO^* brains 48 hours after t-MCAO. Each dot represents an animal (n=8; data are means ± s.e.m.; n.s. p>0.05 (not shown); Student’s t-test). (**F**) Quantification of serum IgG leakage area in each bregma region of WT and *Rab7a^iECKO^*brains 48 hours after t-MCAO (Each dot represents the average of n=8 mice / group; data are means ± s.e.m. one-way ANOVA with post-hoc Tukey’s correction). (**G**) Quantification of IgG leakage intensity in the ipsilateral cortex of each bregma region in WT and *Rab7a^iECKO^* brains 48 hours after t-MCAO. The dotted line represents the average fluorescence intensity in the contralateral cortex (Each dot represents the average of n=8 mice / group; data are means ± s.e.m.; one-way ANOVA with post-hoc Tukey’s correction). (**H**) Quantification of 70 kDa-Dextran-TMR leakage in WT and *Rab7a^iECKO^* ipsilateral and contralateral cortices 48 hours after t-MCAO. Each dot represents an animal [n=2 healthy mice, n=4 mice / genotype for stroke; data are means ± s.e.m.; n.s. p>0.05 (not shown); Mann-Whitney t-test]. (**I**) Western blot for Caveolin-1 protein levels in lysates collected from the contralateral and ipsilateral cortices of *WT* and *Rab7a^iECKO^* brains 48 hours after t-MCAO. (**J**) Quantification of Caveolin-1 protein levels in contralateral and ipsilateral cortical lysates from *WT* and *Rab7a^iECKO^* brains 48 hours after t-MCAO, normalized to contralateral WT levels. Each dot represents an animal (n=6 mice / group; data are means ± s.e.m.; *: p<0.05; n.s.: p>0.05 (not shown); one-way ANOVA with post-hoc Tukey’s correction).

### *Rab7a^iECKO^* mice show improved neuronal outcomes 48 hours after t-MCAO

To determine whether Rab7a-mediated rescue in acute paracellular BBB permeability affect neuronal health after t-MCAO, we assessed the quality and number of NeuN^+^ neurons in the ipsilateral (penumbra) and contralateral somatosensory cortices of both genotypes at seven distinct bregmas. NeuN staining appeared missing or fragmented in the ipsilateral somatosensory cortex of *Rab7a^fl/fl^* (WT) mice, but it appeared largely normal in the ipsilateral cortex of *Rab7a^iECKO^* mice (**Figure 4A-I’**). Quantification of the ratio of aberrant NeuN^+^ cells / DAPI^+^ cells showed a significant 2-fold reduction in the ipsilateral compared to the contralateral cortex of WT mice (**Figure 4J**). However, the ratio of aberrant-appearing NeuN^+^ cells / DAPI^+^ cells was comparable between the ipsilateral and contralateral cortexes of *Rab7a^iECKO^* mice in all bregma regions (**Figure 4J**). We could not detect any cleaved-Caspase3^+^ apoptotic neurons in the stroke region at 48 hours post-t-MCAO (data not shown). Consistent with a greater number of normal appearing cortical NeuN^+^ neurons, *Rab7a^iECKO^*showed fewer neurological deficits with a lower modified Bederson neurological score (0-5) using a series of behavioral tests (forelimb flexion, circling, resistance to lateral push as published ^44^), compared to WT mice by 48 hours after t-MCAO (**Figure 4K**). Therefore, ablation of endothelial Rab7 rescues both acute BBB disruption and neuronal injury in the penumbra region at 48 hours post-t-MCAO.

**Figure 4.**
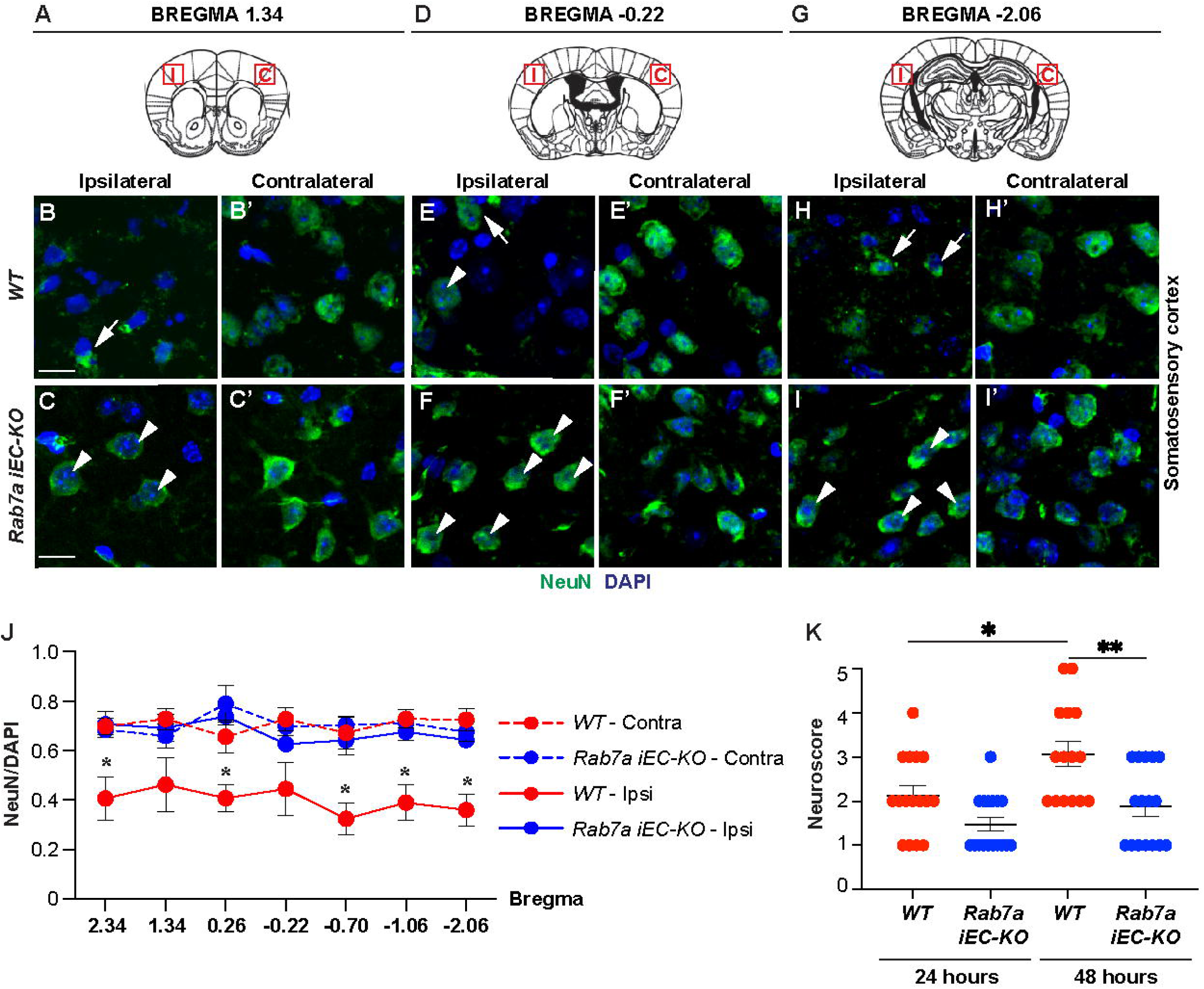
*Rab7a^iECKO^* mice show improved neuronal survival and neurological score 48 hours after t-MCAO. (**A-I’**) Immunofluorescence analysis for NeuN (green) and DAPI (blue) in the contralateral and ipsilateral cortex of WT and *Rab7a^iECKO^* mice 48 hours after t-MCAO. (**A, D, G**) Diagrams illustrate the bregma regions and the red boxes outline the cortical area shown in each micrograph in **B-I**’. (**B-I’**) White arrows indicate abnormal and white arrowheads indicate normal NeuN staining in the ipsilateral cortex. Scale bar: 25 μm. (**J**) Quantification of the fraction of viable neurons (NeuN/DAPI ratio) in the contralateral (dotted lines) and ipsilateral (solid lines) cortices in seven bregma regions of WT and *Rab7a^iECKO^* mice 48 hours after t-MCAO. Each dot represents the average of n=8 animals / group. Data are means ± s.e.m. *: p<0.05; one-way ANOVA with post-hoc Tukey’s correction. (**K**) Analysis of neurological score in WT and *Rab7a^iECKO^* mice 24 and 48 hours after t-MCAO. Each dot represents one animal (n=14 WT and n=16 *Rab7a-iECKO* mice). Data are medians ± 95% C.I. *: p<0.05; **: p<0.01; Kruskal-Wallis one-way ANOVA.

To examine if the reduced paracellular BBB permeability found in *Rab7a^iECKO^* mice after ischemic stroke is accompanied by reduced inflammatory responses of glial cells, we analyzed the percentage of activated myeloid cells (activated microglia and infiltrating macrophages) and resting microglia in the stroke core and border regions of the ipsilateral cortex and contralateral cortex of both genotypes 48 hours after t-MCAO in three bregmas with significant BBB damage rescue (**Figure 5A-I**). Resting myeloid cells (mostly Iba1^+^ CD68^-^ microglial cells with ramified branches) were found outside the BBB leakage area in the border region of the ipsilateral cortex and contralateral cortex (**Figure 5A-B’”, D-E’”**; open arrowheads). In contrast, activated myeloid cells (Iba1^+^ CD68^+^ cells with ameboid morphology) were found in the core and border regions of the ipsilateral cortex in both WT and *Rab7a^iECKO^* cortexes (**Figure 5B-C’”, E-F’”**; yellow arrows). Although the ratio of CD68^+^ Iba1^+^ over total Iba1^+^ cells was higher in the core and border regions in the ipsilateral cortex compared to the contralateral cortex, there were no significant differences between the two genotypes (**Figure 5G-I**). In addition, there was no difference in the mean fluorescence intensity of GFAP reactivity, a marker of activated astrocytes, in the border ipsilateral regions in both genotypes at 48 hours post-t-MCAO (**Figure 5J-L**). The comparable inflammatory responses of both myeloid cells and astrocytes between WT and *Rab7a^iECKO^* cortexes in the acute phase (48 hours) of ischemic stroke could be a consequence of a partial, rather than a complete, rescue in BBB permeability in *Rab7a^iECKO^* mice.

**Figure 5.**
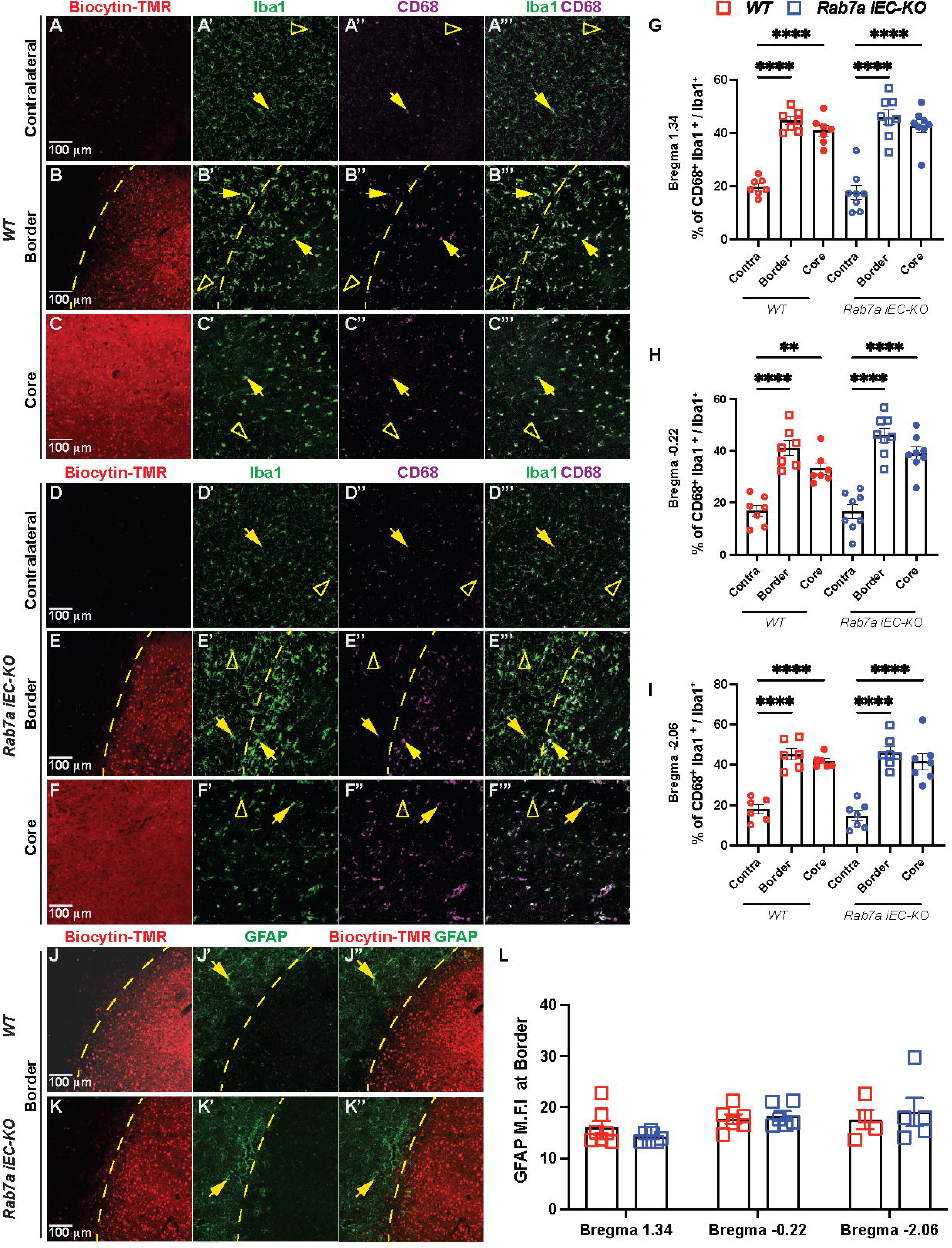
Rab7a-mediated rescue of BBB permeability after t-MCAO is not associated with reduced microglia activation / macrophage infiltration or astrocyte activation. **(A-F’’’)** Immunofluorescences images for biocytin-TMR (red), Iba1 (green), and CD68 (magenta) in the contralateral cortex **(A-A’’’,D-D’’’)**, border **(B-B’’’, E-E’’’)**, and core **(C-C’’’,F-F’’’)** regions of the ipsilateral cortex of *Rab7a^fl/fl^ (WT)* and *Rab7a^iECKO^*mice 48 hours after t-MCAO. Yellow arrowheads point to Iba1^+^, CD68^+^ activated myeloid cells, open yellow arrowhead indicate Iba1^+^ resting microglia. Scale bar = 100 μm. Quantification of activated / resting myeloid cells at bregmas **(G)** 1.34, **(H)** −0.22, and **(I)** −2.06 in the contralateral cortex, border and core regions of the ipsilateral cortex of *WT* and *Rab7a^iECKO^*mice 48 hours after t-MCAO. Each dot represents an animal (n= 6-9 mice/group). Data are means ± s.e.m. ***: p<0.001; **: p<0.01; *: p<0.05; one-way ANOVA with post-hoc Tukey’s correction. **(J-K’’)** Immunofluorescences images for biocytin-TMR (red), and GFAP (green) in the border of the ipsilateral cortex of *WT* and *Rab7a^iECKO^* mice 48 hours after t-MCAO. Yellow arrowheads point to GFAP^+^ reactive astrocytes at the stroke border. Scale bar = 100 μm. **(L)** Quantification of GFAP mean florescence intensity (M.F.I) at the stroke border across three bregmas 1.34, −0.22, −2.06 in *WT* and *Rab7a^iECKO^* mice 48 hours after t-MCAO. Data are means ± s.e.m and normalized to WT mice. ***: p<0.001; **: p<0.01; *: p<0.05; one-way ANOVA with post-hoc Tukey’s correction.

### Endothelial Rab7a elimination reduces structural abnormalities at the BBB tight junctions 48 hours after t-MCAO

To understand how Rab7a affects the disassembly of BBB tight and adherens junctions during the acute phase of ischemic stroke, we analyzed the morphology of eGFP::Claudin-5^+^, ZO-1^+^ TJs and CDH5^+^ AJs at 48 hours after t-MCAO by immunofluorescence and electron microscopy. We first quantified the fraction of Glut1^+^ vessel segments with either intact eGFP^+^ junction strands, junction strands with large gaps (> 2.5 μm), or absent junctions from 6-8 independent fields (ROIs) located at the core, border and contralateral cortical areas (**Figures 6A-C”, 6D-F”**) of two bregma regions (0.26 to −0.22) with the highest rescue in acute BBB leakage (**Figure 3F**). In the last category, we also included vessel segments where eGFP::Claudin-5^+^ protein was located inside the BECs (**Figure 6F”**). The majority (~63-66%) of eGFP::Claudin-5^+^ TJs strands were intact in the contralateral regions regardless of the genotype, and this fraction was significantly decreased (~30.6%) in the core region of *Rab7a^fl/fl^* (WT) mice at 48 hours after t-MCAO (**Figure 6A-A”’, G**). In contrast, there was a significantly higher fraction of vessel segments with TJ strand gaps [~50%; **Figure 6B-B”’, H**] and absent junctions or intracellular GFP^+^ inclusions [~20%; **Figure 6C-C”’, I**] in the core, compared to the border and contralateral regions of WT mice. Although, the fraction of vessel segments with TJ strand gaps was reduced in the core region of *Rab7a^iECKO^*cortices (~40%) and there was no longer a significant difference among the ipsilateral core, the ipsilateral border and the contralateral regions in *Rab7a^iECKO^* mice for these parameters (TJ strand gaps), we could not detect significant differences either in the fraction of vessel segments with TJ strand abnormalities (**Figure 6E-F”’, H, J**), or vessels area covered by the eGFP^+^ signal (**Figure 6J**) between two genotypes. This finding likely reflects the large variability within the ipsilateral and border regions of the mice within the same group. Ve-Cadherin+ (CDH5^+^) AJs showed a similar behavior to eGFP::Claudin-5^+^ TJs (90-95% correlation) at the BBB after ischemic stroke with no significant differences observed between two genotypes (**Figure S1**). In contrast, *Rab7a^iECKO^* mice had reduced fraction of vessel segments with ZO-1^+^ TJ strands with gaps (~32%) or absent ZO1^+^ TJs (4%) in the ipsilateral cortical core region compared to WT mice [ZO-1^+^ TJ strand with gaps (~49%) and absent TJs (23.2%); **Figure 7A-I**]. Moreover, *Rab7a^iECKO^*mice had a significant higher fraction (65%) of vessels segments with intact ZO-1^+^ TJ strands compared to WT mice (~27.5%) in the ipsilateral cortical core region at 48 h post-t-MCAO (**Figure 7A-A”’, D-D”’, G**). Overall, these findings demonstrate that Rab7a elimination in BECs predominantly reduces ZO-1 degradation and localization to cell junctions compared to Claudin-5 or CDH5 at during the acute BBB damage after ischemic stroke.

**Figure 6.**
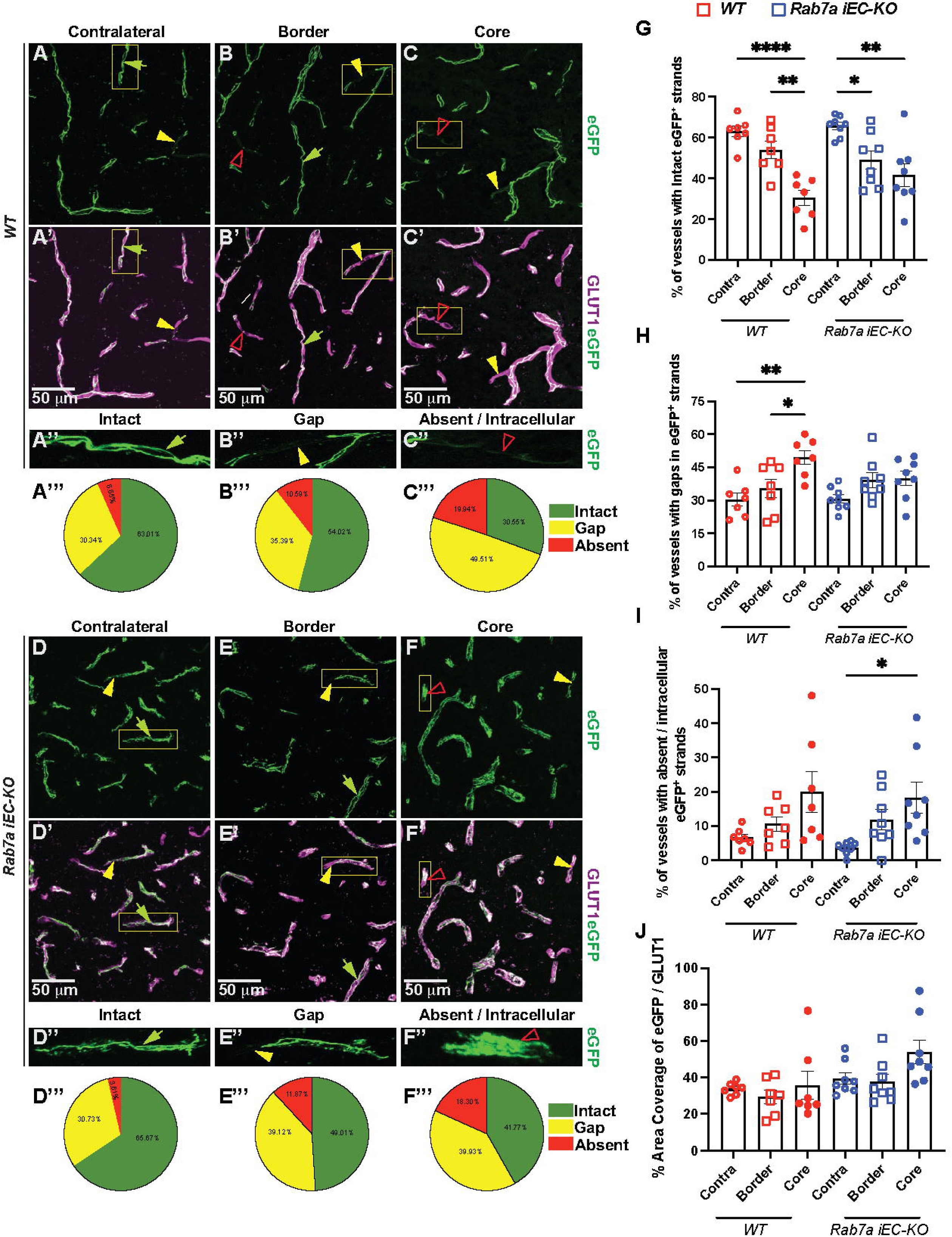
Endothelial Rab7a elimination does not rescue structural abnormalities of eGFP::Claudin-5 tight junctions 48 hours after t-MCAO. (**A-F”’**) Immunofluorescence images for eGFP::Claudin5 (green) and Glut1 (magenta) in the contralateral cortex **(A-A’, D-D’)**, border **(B-B’, E-E’)** and core **(C-C’, F-F’)** regions of the ipsilateral cortex of *WT* and *Rab7a^iECKO^* mice 48 hours after t-MCAO. Normal green arrowhead indicates intact eGFP::Claudin-5^+^ TJ strands, yellow arrowhead points towards gaps in eGFP::Claudin-5^+^ TJ strands, red open arrowheads indicate either absent TJ strands or intracellular eGFP:;Claudin-5^+^ aggregates. Scale bar = 50 µm. **(A”-F”)** Magnified images of the white boxed areas in **(A-F’)** to illustrate the phenotype of the TJ strand. **(A”’-F”’)** Pie charts correspond to the percentage of intact (green), gap (yellow), and absent (red) eGFP::Claudin-5^+^ junctional strands present in the contralateral and ipsilateral cortex of *WT* and *Rab7a^iECKO^* mice 48 hours after t-MCAO. **(G-I)** Dotted bar graphs of the percentage of vessel segments with (**G**) intact TJ segments, **(H)** TJ strands with gaps, **(I)** absent or intracellular eGFP::Claudin-5^+^ aggregates in the contralateral cortex, border and core regions of the ipsilateral cortex of *WT* and *Rab7a^iECKO^* mice 48 hours after t-MCAO. **(J)** Dotted bar graph of the percentage of vascular area covered by eGFP^+^ TJ strands. Each dot represents an animal (n= 7/8 mice/group). Data are means ± s.e.m. ***: p<0.001; **: p<0.01; *: p<0.05; one-way ANOVA with post-hoc Tukey’s correction.

**Figure 7.**
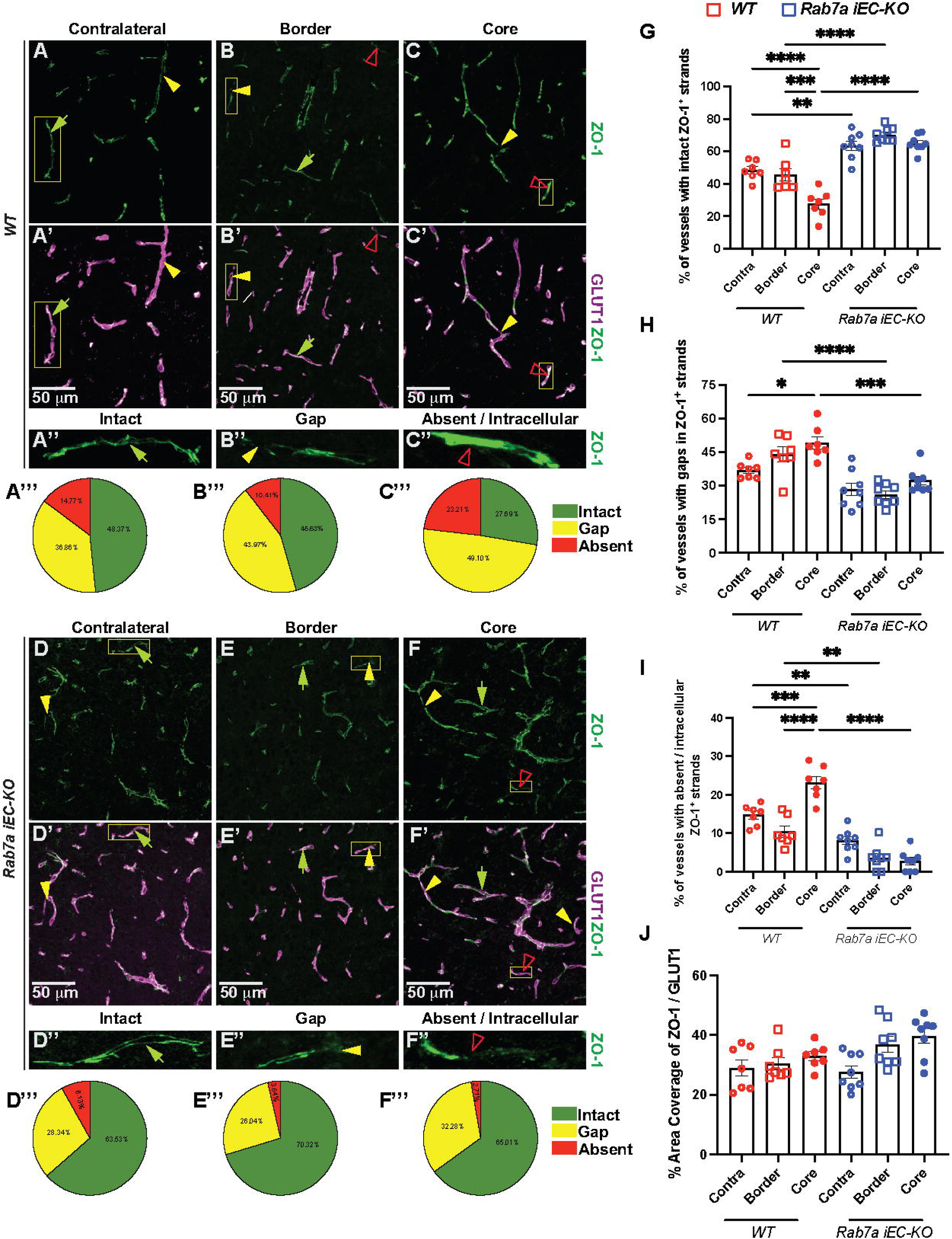
Endothelial Rab7a elimination reduces ZO-1 structural abnormalities in tight junctions 48 hours after t-MCAO. (**A-F”’**) Immunofluorescence images for ZO-1 (green) and Glut1 (magenta) in the contralateral cortex **(A-A’,D-D’)**, border **(B-B’,E-E’)** and core **(C-C’,F-F’)** regions of the ipsilateral cortex of *WT* and *Rab7a^iECKO^* mice 48 hours after t-MCAO. Normal green arrowhead indicates intact ZO-1^+^ TJ strands, yellow arrowhead points towards gaps in ZO-1^+^ TJ strands, red open arrowheads indicate absent ZO-1+ TJ strands. Scale bar = 50 µm. **(A’’-F’’)** Magnified images of the white boxed areas in **(A-F’)** to illustrate the phenotype of the ZO-1^+^ TJ strand. **(A”’-F”’)** Pie charts correspond to the percentage of intact (green), gap (yellow), and absent (red) ZO-1^+^ junctional strands in the contralateral and ipsilateral cortex of *WT* and *Rab7a^iECKO^* mice 48 hours after t-MCAO. **(G-I)** Dotted bar graphs of the percentage of vessel segments with (**G**) intact TJ segments, **(H)** TJ strands with gaps, **(I)** absent ZO-1 in the contralateral cortex, border and core regions of the ipsilateral cortex of *WT* and *Rab7a^iECKO^* mice 48 hours after t-MCAO. **(J)** Dotted bar graph of the percentage of vascular area covered by ZO-1^+^ TJ strands. Each dot represents an animal (n= 7/8 mice/group). Data are means ± s.e.m. ***: p<0.001; **: p<0.01; *: p<0.05; one-way ANOVA with post-hoc Tukey’s correction.

Next, we examined the ultrastructural morphology of endothelial TJs in the ipsilateral core cortices of WT and *Rab7a^iECKO^* mice at 48 hours after t-MCAO with transmission electron microscopy. We acquired 16-22 images per cortex and analyzed 46-55 endothelial TJs per animal. At 48 hours after t-MCAO, on average 45% of BBB TJs in the ipsilateral cortex of *Rab7a^fl/f^*^l^ (WT) mice showed structural abnormalities containing large gaps where adjacent membranes were separated from each other due to loss of the electron dense material (**Figure 8A, A’, red arrowhead, C**), consistent with our prior findings in ischemic stroke ^11^. In contrast, the fraction of TJ with these structural abnormalities was significantly reduced (~25%) in *Rab7a^iECKO^* ipsilateral cortex at 48 hours after t-MCAO (**Figure 8B, B’, C**). Thus, Rab7a elimination in BECs reduces acute structural abnormalities at the electron microscopy level of BBB TJs after ischemic stroke.

**Figure 8.**
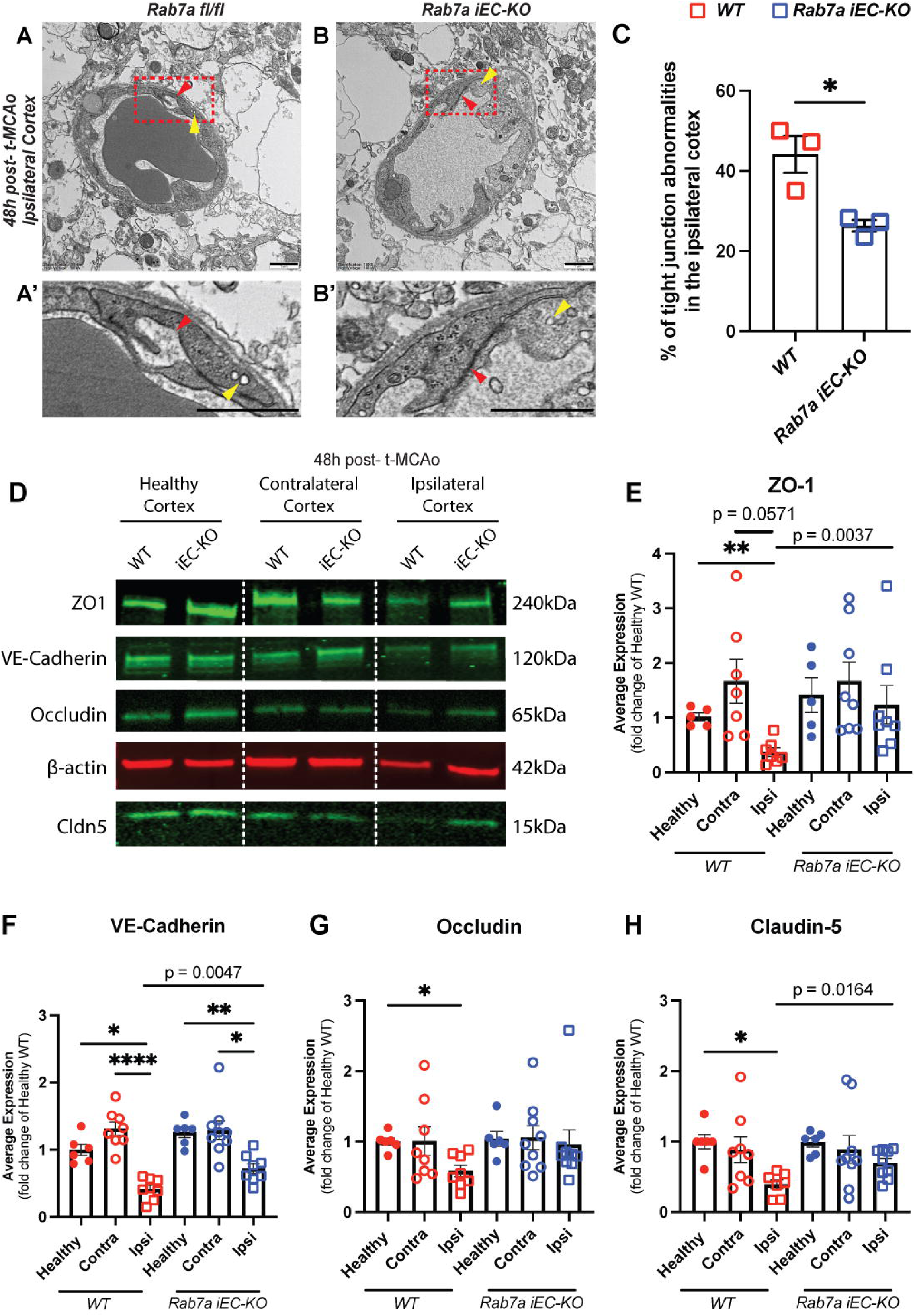
Rab7a regulates degradation of select BBB junctional proteins 48 hours after t-MCAO. (**A-B’**) Transmission Electron Microscopy (TEM) images of tight junctions 48 hours post-t-MCAO in the ipsilateral cortex of *Rab7a^fl/fl^* (WT) and *Rab7a^iECKO^*mice. Red arrowhead point to tight junctions, yellow arrowhead point to caveolae. **(A’-B’)** Magnified images of boxed areas (**A, B**) to illustrate normal and abnormal tight junctions in brain endothelial cells. Scale bar = 1 µm. **(C)** Quantification of tight junction abnormalities in the ipsilateral cortex of WT and *Rab7a^iECKO^* mice. The data were analyzed from 46-55 tight junctions per mouse obtained from 16-22 images in the cortex region (n=3 mice / genotype). Data are means ± s.e.m. *: p<0.05; Mann-Whitney t-test. (**D**) Western blots for BBB junctional proteins ZO-1, VE-cadherin, Occludin and Claudin-5 and β-actin (control) of brain lysates collected from either the healthy, contralateral and ipsilateral ischemic cortex of *WT* and *Rab7a^iECKO^* mice 48 hours after t-MCAO. The respective molecular weights for each protein are shown on the right. (**E-H**) Quantification of Claudin-5, Occludin, ZO-1 and VE-cadherin protein levels in either healthy, contralateral or ipsilateral ischemic brain lysates of *WT* and *Rab7a^iECKO^* mice 48 hours after t-MCAO. Each dot represents an animal (6-9 mice / group). Data are means ± s.e.m and normalized to protein levels in WT healthy mice. ***: p<0.001; **: p<0.01; *: p<0.05; Brown-Forsythe and Welch ANOVA. The p values shown for the comparisons between the ipsilateral ischemic brain lysates of *WT* and *Rab7a^iECKO^* mice 48 hours after t-MCAO were achieved by Mann-Whitney t-test.

To determine the effects of endothelial Rab7a elimination on AJ (VE-Cadherin) and TJ [Claudin-5, Occludin, and ZO-1] protein levels in BECs after ischemic stroke, lysates were collected from the ipsilateral and contralateral cortices of both genotypes at 48 hours after t-MCAO and assessed by Western blotting (**Figure 8D**). VE-Cadherin, Claudin-5, Occludin and ZO-1 protein levels were significantly reduced by approximately 2-fold in the ipsilateral cortex of WT mice at 48 hours after t-MCAO, compared to both the WT healthy (t-MCAO sham) and WT contralateral cortex (**Figure 8E-H**). Similarly, VE-Cadherin protein was significantly reduced in *Rab7a^iECKO^* ipsilateral cortex compared to *Rab7a^iECKO^* healthy and contralateral cortex (**Figure 8F**). In contrast, there was no significant difference in Claudin5, Occludin or ZO-1 in *Rab7a^iECKO^*ipsilateral cortex compared to *Rab7a^iECKO^* healthy and contralateral cortex when we performed a group analysis (Brown-Forsythe Welsh ANOVA) in this cohort of mice [ZO1 (p =0.3479), VE-Cadherin (p = 0.0694), and Claudin5 (p=0.0717)] likely due to smaller effect size, the larger variability within the same group and the relatively smaller number of mice (data not shown). However, when we compare the AJ and TJ protein levels between the ipsilateral cortexes of the two genotypes only using a Mann-Whitney t-test, there is a significant increase in ZO1 (p =0.0037), VE-Cadherin (p = 0.0047), and Claudin5 (p=0.0164) levels in the ipsilateral *Rab7^iECKO^* compared to WT cortex (**Figure 8E, G, H**). These data support the idea that Rab7a elimination in BECs can partially rescue the degradation of select TJ or AJ proteins after ischemic stroke, although the effects are not very robust.

### Pro-inflammatory cytokines, TNF**α** and IL1**β**, but not glucose and oxygen deprivation, induce Rab7a activation in BECs *in vitro*

Rab7a cycles between inactive (GDP-bound) and active (GTP-bound) states to regulate various cell biological processes ^28, 29^. Rab-interacting lysosomal protein (RILP) binds selectively to Rab7-GTP and recruits the dynein-dynactin motor complex to facilitate vesicle movement toward the minus end of microtubules ^45^. The N terminus region of RILP is required to recruit dynein motors, but not for Rab7-GTP binding; therefore an N-terminal truncation of RILP can be used to assess Rab7 activation in cells ^33^. To investigate which cytokines may induce Rab7a activation in ischemic stroke, we cultured primary mouse BECs *in vitro* under conditions of either glucose and oxygen deprivation (OGD), or inflammation (TNFα and IL-1β; 10 ng/mL) for 48 hours and quantified the amount of GTP-bound Rab7a by using a GST-RILP fusion protein to pull down selectively the active protein from cell lysates (**Figure 9A**). Both the total and active Rab7a levels were decreased significantly after 48 hours of OGD treatment (**Figure 9G-I**), suggesting that ischemic conditions do not activate Rab7a protein. Since ischemic stroke induces inflammation which furthers participates in BBB disruption^14^, we then tested whether distinct pro-inflammatory cytokines induce Rab7a activation. Treatment with TNFα and IL1β did not increase the total amount of Rab7a, but increased, albeit non-significantly, the GTP-bound Rab7a levels after 24 hours (**Figure 9B-D**). After 48 hours of TNFα and IL1β treatment, Rab7a-GTP levels, but not total Rab7a levels, were significantly increased (**Figure 9B, E, F**). Three other pro-inflammatory cytokines that are upregulated after ischemic stroke, IL-21 ^46^, CCL2 ^47^ and IL-17A ^48^, had no effect on either total or activated Rab7a levels in BECs (**Figure S2**). Therefore, select pro-inflammatory cytokines promote Rab7a activation in primary BECs. Next, we examined expression of Ccz1, a GEF that acts together with vacuolar fusion protein Mon1 to promote both Rab7a activation and Rab7a-mediated membrane fusion in yeast ^35^ and eukaryotic cells ^36–39^. Ccz1 expression was increased, albeit non-significantly, in BECs 24 to 48 hours after TNFα and IL1β exposure (**Figure S2J, K**). Thus, TNFα and IL1β may upregulate expression of Ccz1 protein in BECs within 24 hours, which could activate Rab7a to promote degradation of AJ and TJ proteins.

**Figure 9.**
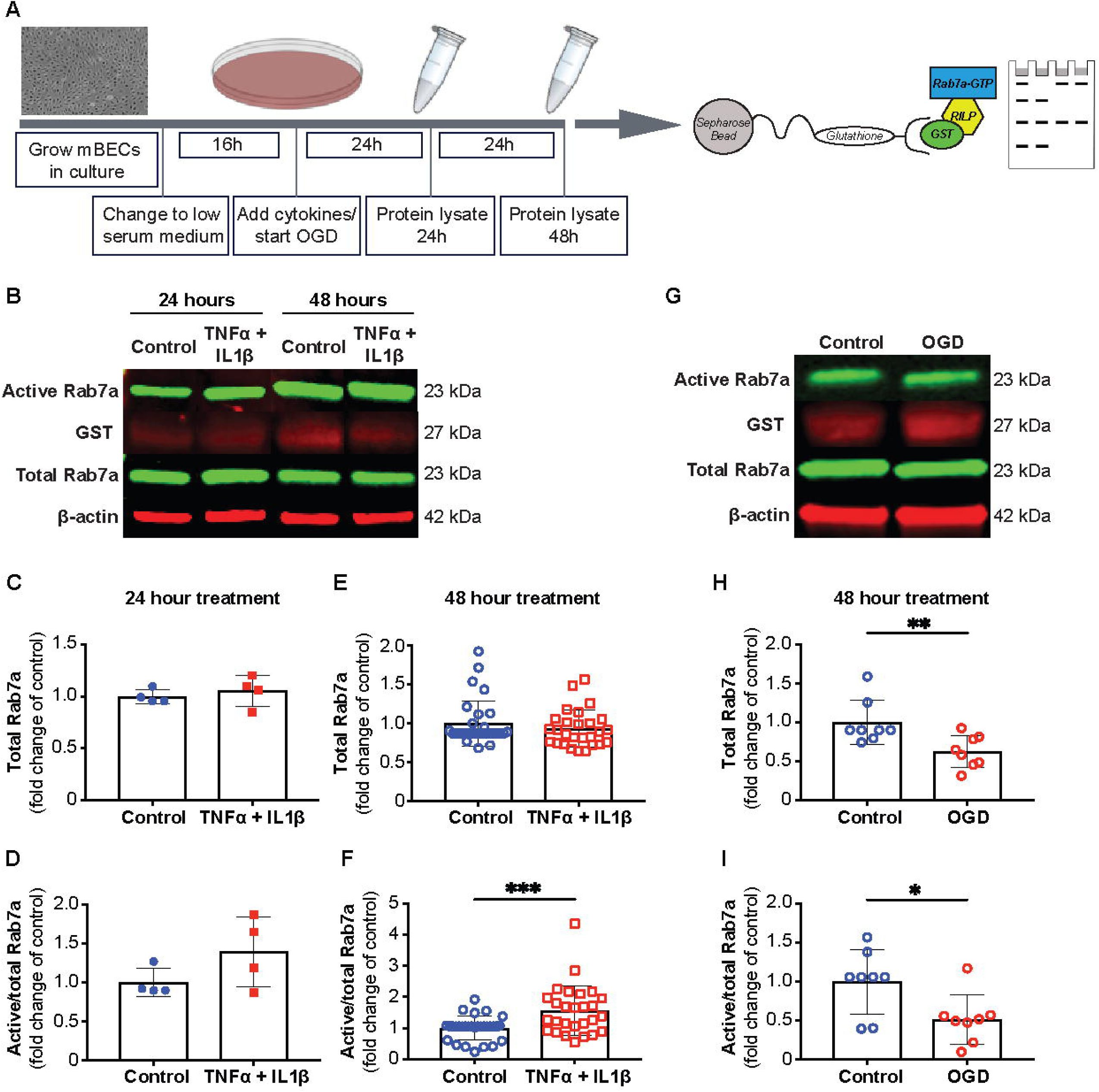
Proinflammatory cytokines TNFα and IL1β induce Rab7a activation in mouse brain endothelial cells. (**A**) Schematic diagram of the experimental setup. Primary mouse brain endothelial cells (mBECs) are grown to confluence and switched to low-serum media overnight prior to addition of pro-inflammatory cytokines (TNFα, IL-1β; 10 ng/mL). Protein lysates are collected after 24 or 48 hours of cytokine treatment. Rab7a-GTP is pulled down using a GST-RILP fusion protein that binds glutathione immobilized on sepharose beads. Western blotting is then used to detect both Rab7a and Rab7a-GTP protein levels. (**B**) Western blot of active and total Rab7a proteins in mBECs treated with TNFα and IL1β for either 24 or 48 hours and untreated control cells. GST and β-actin are used to normalize active Rab7a and total Rab7a levels, respectively. The molecular weight of each protein is shown on the right. (**C-F**) Quantification of total (**C, E**) and active (**D, F**) Rab7a levels in mBECs treated with TNFα and IL1β for 24 hours (**C, D**) and 48 hours (**E, F**) and untreated cells. Each dot represents an independent experiment. Data are means ± s.e.m. (**F**) ***: p<0.005; Student’s t-test. (**G**) Western blot for active and total Rab7a in mBECs grown in oxygen and glucose deprivation (OGD) conditions for 48 hours and untreated control cells. GST and β-actin are used to normalize active Rab7a and total Rab7a levels, respectively. The molecular weight of each protein is shown on the right. (**H, I**) Quantification of total (**H**) and active (**I**) Rab7a levels in mBECs grown in OGD conditions for 48 hours and control cells. Each dot represents an independent experiment. Data are means ± s.e.m.; **: p<0.01; *: p<0.05; n.s.: p>0.05 (not shown) Student’s t-test.

### Silencing Rab7a *in vitro* rescues partially cytokine-driven barrier disruption by reducing internalization of some junctional proteins and formation of F-actin bundles at cell junctions

To investigate how Rab7a affects BEC barrier integrity at the cell biological level, we generated Rab7a knockdown primary mouse BECs via transfection of stealth RNAi duplexes targeting the *Rab7a* mRNA. Rab7a protein levels were decreased by more than 95% in BECs transfected with the siRNA (siRab7a) compared to those transfected with a scrambled siRNA (siCTRL) by immunofluorescence and Western blotting. This knockdown efficiency was observed in the presence and absence of TNFα and IL1β (**Figure S3A-F**). Rab7a knockdown did not affect the levels of several endolysosomal pathway proteins including the early endosome marker EEA-1 and the lysosomal marker LAMP-1 as they were similar in both siCTRL- and siRab7a-transfected mBECs with or without cytokine treatment (**Figure S3E, G, I**). However, Rab7a knockdown increased Ccz1 protein levels in mBECs compared to the siCTRL group without cytokine treatment indicating that Rab7a may be a target for Ccz1. Ccz1 levels were not different at 48 hours after cytokine treatment between the two groups (**Figure S3E, H**). Since Rab7a activity maintains lysosome mass, we quantified the lysosome mass per cell in both siCTRL- and siRab7a-transfected mBECs after immunostaining for the pH sensor Lysotracker. The lysosome mass per cell was similar between siCTRL and siRab7a treatments with or without cytokine treatment (**Figure S3J-N**). Thus, Rab7 knockdown does not affect either other endolysosomal proteins, or the lysosome mass in mBECs.

Since Rab7a plays a critical role in autophagy (reviewed in ^49^), we examined whether Rab7a knockdown affected expression of the two autophagy proteins Atg5 and p62 under basal and inflammatory conditions in mBECs. Under basal conditions, Atg5 and p62 protein levels were similar in both siCTRL- and siRab7a-transfected mBECs (**Figure S3O-Q**). As expected, the Atg5 and p62 protein levels were upregulated under inflammatory conditions in both siCTRL- and siRab7a-transfected mBECs (**Figure S3O-Q**). Moreover, p62 levels were significantly higher in siRab7a-compared to siCTRL-transfected mBECs after cytokine treatment, suggesting that Rab7a knockdown induced p62 accumulation in mBECs (**Figure S3O-Q**), likely due to due to increased cellular stress or impaired autophagic reflux.

To determine if Rab7a knockdown rescues cytokine-driven increase in mouse BEC paracellular permeability we analyzed the transendothelial electrical resistance (TEER) in siCTRL- and siRab7a-transfected cells after cytokine treatment. In the absence of cytokines, siRab7a-transfected mBECs show a decreased basal TEER compared to siCTRL-transfected cells (**Figure S4A-C**). Treatment of siCTRL-transfected mBECs with TNFα and IL1β for 48 hours significantly reduced the TEER by 30%. However, this cytokine mediated TEER reduction was less effective in siRab7a-transfected BECs (15%; **Figure S4A-C**). To investigate which cell junction proteins are more sensitive to Rab7a-mediated degradation in BECs, we examined total levels and subcellular localization of AJs (VE-Cadherin) and TJ proteins (Claudin-5, Occludin and ZO-1) at 48 hours after cytokine addition. TNFα and IL1β treatment significantly reduced Claudin-5 and ZO-1, but not VE-Cadherin or Occludin, protein levels between 2- to 3-fold in siCTRL-transfected mBECs (**Figure S4D-H**). siRab7a knockdown did not show a significant rescue in cytokine-induced degradation of VE-Cadherin, ZO-1, Occludin or Claudin-5 proteins in mBECs by LICOR western blotting, suggesting that the cytokine impact on AJ and TJ total protein levels is largely independent of Rab7a *in vitro* (**Figure S4D-H**). By immunofluorescence, siRab7a-transfected mBECs showed normal expression of Claudin-5, ZO-1, VE-Cadherin and α-Catenin (AJs) at cell junctions similar to siCTRL-transfected BECs, although Claudin-5 and VE-Cadherin proteins could also be detected inside the cell (**Figure S4I, N, K, P)**. Consistent with these data, the analysis of VE-cadherin internalization with the mCling assay was increased in siRab7a-transfected compared to siCTRL-transfected mBECs (**Figure S5H-K”**). Treatment with TNFα and IL1β triggered loss of Claudin-5 and VE-Cadherin at cell junction and increased intracellular aggregates in siCTRL-treated BECs (**Figure S4J, K’**). These cytokine-mediated effects were less severe in Rab7a knockdown mBECs where we could observe some Claudin-5 and VE-cadherin proteins still present at cell junctions in addition to intracellular aggregates (**Figure S4O, P’**). However, cytokine treatment increased the rate of VE-Cadherin internalization more in siCTRL than siRab7-transfected mBECs (**Figure S5J-L”**). Cytokine treatment also reduced ZO-1 and α-Catenin junctional localization in a similar manner in control and siRab7a-transfected mBECs (**Figure S4I’, J’, N’, O’, L, L’, Q, Q’**), indicating that Rab7a knockdown affects select AJ and TJ proteins. Moreover, F-actin levels at cell junctions visualized by phalloidin staining were increased significantly in siCTRL-transfected mBECs after TNFα and IL1β treatment, but to a lesser extent in siRab7a-transfected mBECs (**Figure S4M, M’, R, R’**). Since F-actin levels correlate with tension at cell junctions, these data suggest that Rab7a knockdown preserves some of the AJ and TJ proteins at cell junctions following cytokine treatment.

Cav-1 protein can regulate Claudin-5 protein turnover in mBECs after exposure to inflammatory chemokines (e.g. CCL2)^22^, prompting us to examine its levels and the caveolar-mediated transport in Rab7a knockdown BECs. Cav-1 protein levels were increased in primary BECs after exposure to TNFα and IL1β independently of Rab7a levels (**Figure S5F, G**). In addition, we performed an albumin-594 uptake assay to measure Caveolin-1-dependent transcellular uptake, and confirmed that albumin-594 uptake was equally increased in siCTRL- and siRab7a-transfected mBECs upon treatment with cytokines (**Figure S5A-E**). Therefore, the partial rescue of Claudin-5 junctional localization after pro-inflammatory cytokine treatment of siRab7a-transfected mBECs, is not due to changes in Caveolin-1 protein levels and caveolar-mediated uptake and transport. In conclusion, the *in vivo* and some of the *in vitro* data support the hypothesis that Rab7a is necessary in part for cytokine-induced degradation of select AJ and TJ proteins during the acute BBB damage after ischemic stroke.

## DISCUSSION

The disassembly of cell junctions due to persistent degradation or aberrant localization of adherens and tight junction proteins contributes together with other mechanisms to increase BBB permeability in multiple neuropathological conditions, including ischemic stroke (reviewed in ^3^). However, the cell biological mechanisms driving junctional disassembly are not fully understood. OGD has been proposed to promote junctional protein degradation through activation of several MMPs that cleave the extracellular domains of select AJ and TJ proteins in BECs ^50, 51^. Surprisingly, OGD does not activate Rab7a in primary BECs, suggesting that OGD-induced junctional protein degradation does not require Rab7a. On the contrary, IL-1β and TNF-α, which are upregulated in stroke tissue between 24 to 48 hours after reperfusion and coincide with immune cell infiltration into the CNS (reviewed in ^13, 14^), induce internalization and degradation of select BBB AJ and TJ proteins ^22, 26^. One of the mechanisms by which IL-1β and TNF-α mediate BBB cell junction protein degradation is via activation of Rab7a activation, which then degrades select BBB cell junction-associated proteins. Other pro-inflammatory cytokines/chemokines that have been implicated in BBB dysfunction such as IL-17A, IL-21 and CCL2 may promote TJ degradation through Rab7a-independent mechanisms, since they cannot activate Rab7a in primary BECs. Therefore, pro-inflammatory cytokines likely mediate BBB TJ degradation at the acute phase of ischemic stroke through both Rab7a-dependent and -independent mechanisms.

Cav-1 has been previously implicated in tight junction transmembrane protein internalization and degradation ^22, 26, 50^. Our data demonstrate that Rab7a-mediated rescue of acute BBB dysfunction after stroke *in vivo* is not accompanied by changes in Caveolin-1 protein levels or caveolar-mediated transcellular permeability *in vivo* and *in vitro*, although the inflammatory milieu in stroke increases Cav-1 protein levels in BECs. Moreover, the pro-inflammatory chemokine CCL2 does not induce Rab7a activation in primary mBECs, although it has been shown to promote Caveolin-1-mediated internalization of Claudin-5 in the bEnd3.0 cell line ^22^. Although we cannot exclude the possibility that Caveolin-1-dependent cargo sorting is a putative mechanism of select BBB junction protein degradation under some conditions, it is likely not the predominant mechanism by which inflammatory cytokines and chemokines mediate degradation of cell junction proteins in acute ischemic stroke. Thus, the cellular mechanisms of BBB AJ and TJ protein degradation are context dependent and likely differ across distinct neurological diseases and stages of disease.

Overall our findings, primarily *in vivo*, suggest a model for how the state of Rab7a activation in BECs regulates, in part, degradation of select BBB junctional proteins in healthy and disease states (**Figure S5M, N**). We propose that under physiological conditions, similar levels of active and inactive Rab7a protein levels maintain a constant internalization, recycling and degradation of adherens and tight junction proteins, in order to retain constant levels at cell junctions (**Figure S5M**). When BECs are exposed to a subset of pro-inflammatory cytokines (e.g. TNFα and IL1β), such as in acute ischemic stroke, Rab7a is activated, although the mechanism remains unclear. Rab7a may be activated through a Ccz1-mediated mechanism ^35–37^, although we could not test this hypothesis because we could not knockdown Ccz1 in primary mBECs (data not shown). Alternatively, Rab7a may be activated in BECs by reducing TBC1D15 (GAP protein) levels as found *in vitro*^52, 53^ and in a mouse model of myocardial infarction^54^, or through post-translational protein modifications (reviewed in ^55^). Rab7a activation pushes the endolysosomal pathway towards degradation, rather than recycling, and drives cell junctional proteins away from cell junctions toward the lysosomes which further exacerbates acute changes in paracellular BBB permeability after ischemic stroke (**Figure S5N**). Our *in vivo* data suggest that endothelial Rab7 elimination is more effective in reducing abnormal ZO-1 localization at BBB cell junctions after ischemic stroke compared to eGFP::Claudin-5 and VE-Cadherin proteins and preserving their total protein levels, although the effects are not very robust. Overall, our study identifies a new molecular mechanism controlling BEC junctional integrity in diseased BBB, and establishes Rab7a as an important regulator of AJ and TJ protein turnover via the endolysosomal pathway under inflammatory conditions. Our findings have implications not only for ischemic stroke, but also other neurological disorders characterized by inflammation and BBB dysfunction. For example, in EAE, structural abnormalities in BBB TJs precede the onset of disease and persist throughout the course of EAE to allow immune cell infiltration into the CNS ^12^, emphasizing a critical role for pro-inflammatory mechanisms in driving TJ strand disassembly. Based on our findings, we predict that early and persistent Rab7a activation due to pro-inflammatory cytokines Il-1β and TNF-α may likely drive degradation of BBB TJ transmembrane proteins in EAE leading to BBB dysfunction ^12^.

The mechanism by which Rab7a activation triggers degradation of BBB junctional proteins to promote dismantlement of BEC cell junctions after ischemic stroke implies a potential lack of selectivity, since the endolysosomal degradation is a fundamental cell biological process. The genetic loss-of-function data suggest that Rab7a knockdown has no effect on the levels of Caveolin-1 or Slc2a1 (Glut-1), which are both integral membrane proteins. In contrast, Rab7a elimination *in vivo* rescues the levels and localization of select BBB junctional proteins such as ZO-1 and to a lesser extent Claudin-5 and VE-Cadherin *in vivo*, although these effects are not robust. However, it has very little effect on Occludin which is a member of the MARVEL family of transmembrane proteins. In contrast, our *in vitro* data indicate that Rab7a deletion acutely is not sufficient to prevent degradation of total AJ and TJ protein levels after cytokine treatment, although it rescues to some extent the shift from intracellular to junctional localization by immunofluorescence analysis. Consistent with the effects of Rab7a on regulation of junctional localization of VE-Cadherin and Claudin-5 in BECs are the findings that F-actin stress fibers are more robust in siCTRL-compared to siRab7a-transfected mBECs after exposure to inflammatory cytokines. TNFα, which activates Rab7a, is known to promote the formation of actin stress fibers ^56^ that further induce disassembly and internalization of junctional proteins ^57^. However, Rab7a elimination rescues only partially the degradation of ZO-1, Claudin-5 and VE-Cadherin proteins, suggesting that Rab7a-dependent and Rab7a-independent mechanisms may degrade BBB junctional proteins under inflammatory conditions. For example, inflammatory cytokines also promote ubiquitin-mediated proteasome degradation of VE-cadherin ^58^, Claudin-5 ^59^ and ZO-1 (reviewed in ^20^). Rab7a activation is also critical to promote autophagy which together with the ubiquitin-proteasome system serve as two independent mechanisms to degrade and clear protein debris under cellular stress responses such as ischemic stroke (reviewed in ^60^). It is conceivable that both mechanisms are activated by inflammatory cytokines in BECs after ischemic stroke to degrade damaged AJ and TJ proteins and promote formation of new junctions at the BBB for post-stroke recovery. In conclusion, our findings provide evidence for a new mechanism by which inflammatory cytokines promote degradation of select junctional proteins via Rab7a activation, leading to BBB dysfunction after ischemic stroke.

## MATERIALS AND METHODS

### Mice

All experimental procedures were approved by the IACUC committees at the University of California, Irvine and Columbia University Irving Medical Center. The following mouse strains were used: *Tg(eGFP-Claudin5)* ^11^; *Rab7a^fl/fl^* ^34^; *Cdh5(PAC)-Cre^ERT2^ (VEC-PAC)* ^61^. To induce Cre^ERT2^-mediated recombination, 4-OH-tamoxifen (Sigma-Aldrich) was dissolved in 10% ethanol/corn oil mixture to a final concentration of 2 mg/ml, and administered by intraperitoneal injections at a dose of 35 μg/g body weight for 5 consecutive days starting at postnatal day P1 - P5 and 5 consecutive days in the adult ending one week before the t-MCAO procedure.

### Reagents

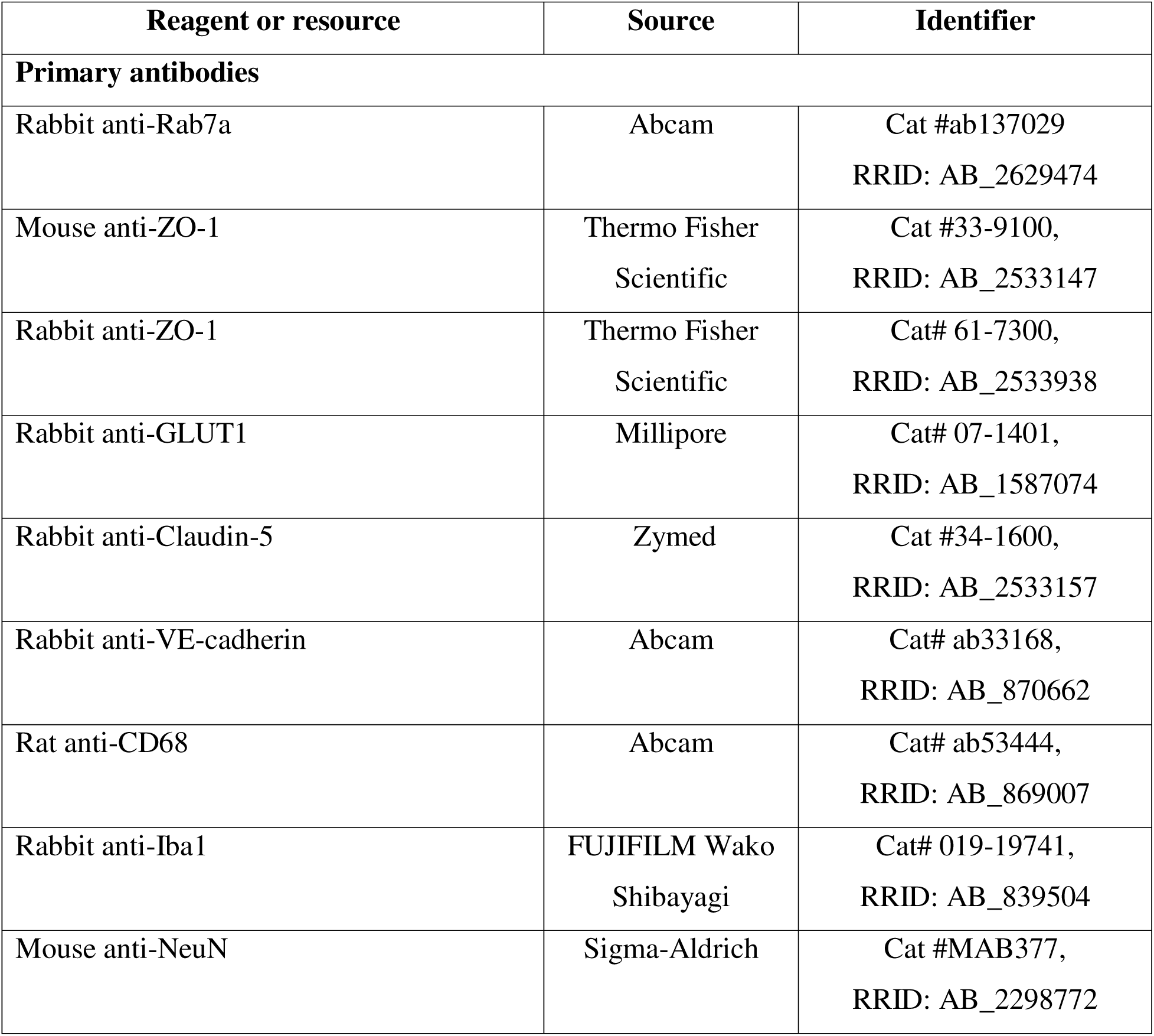
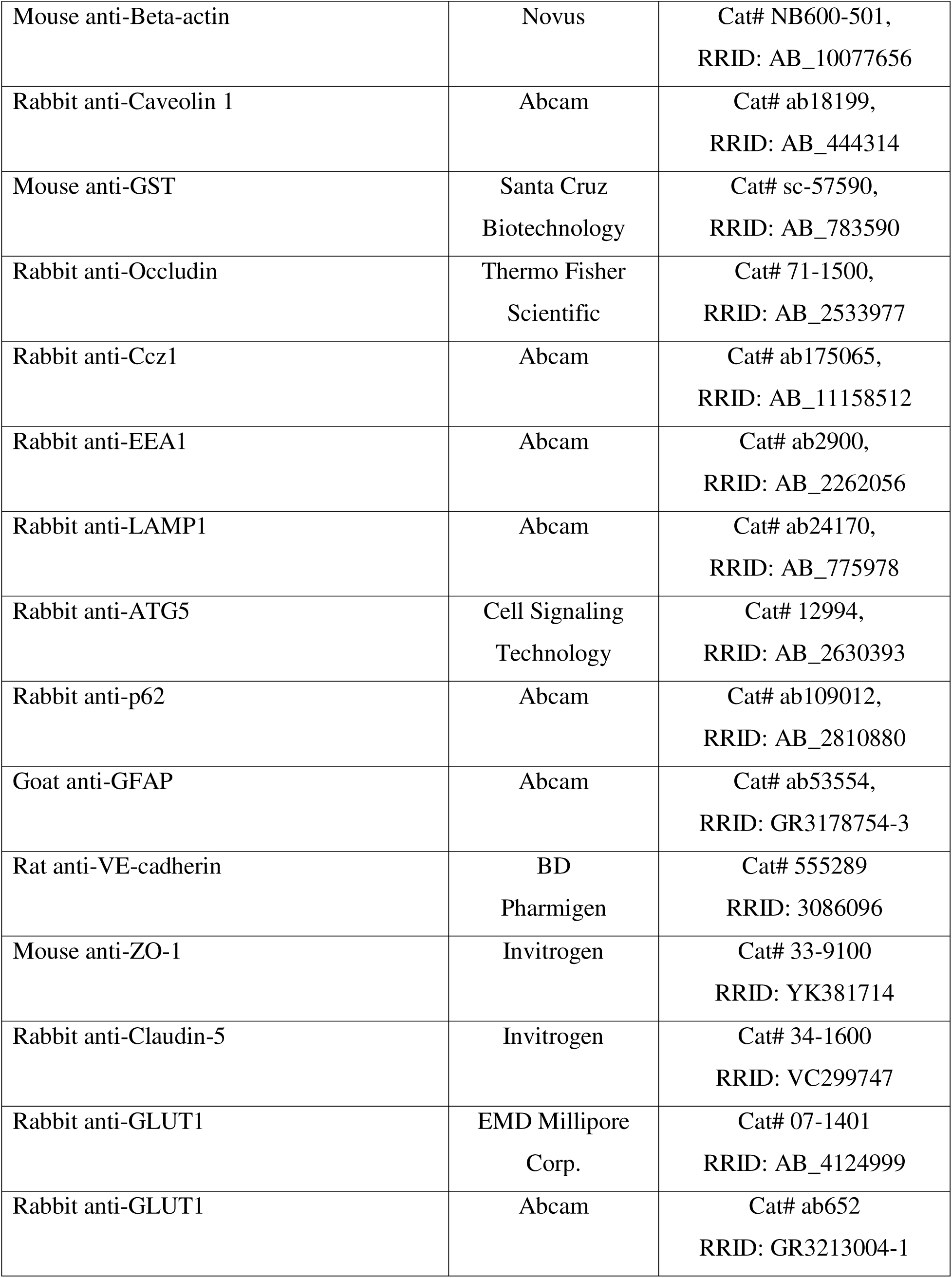
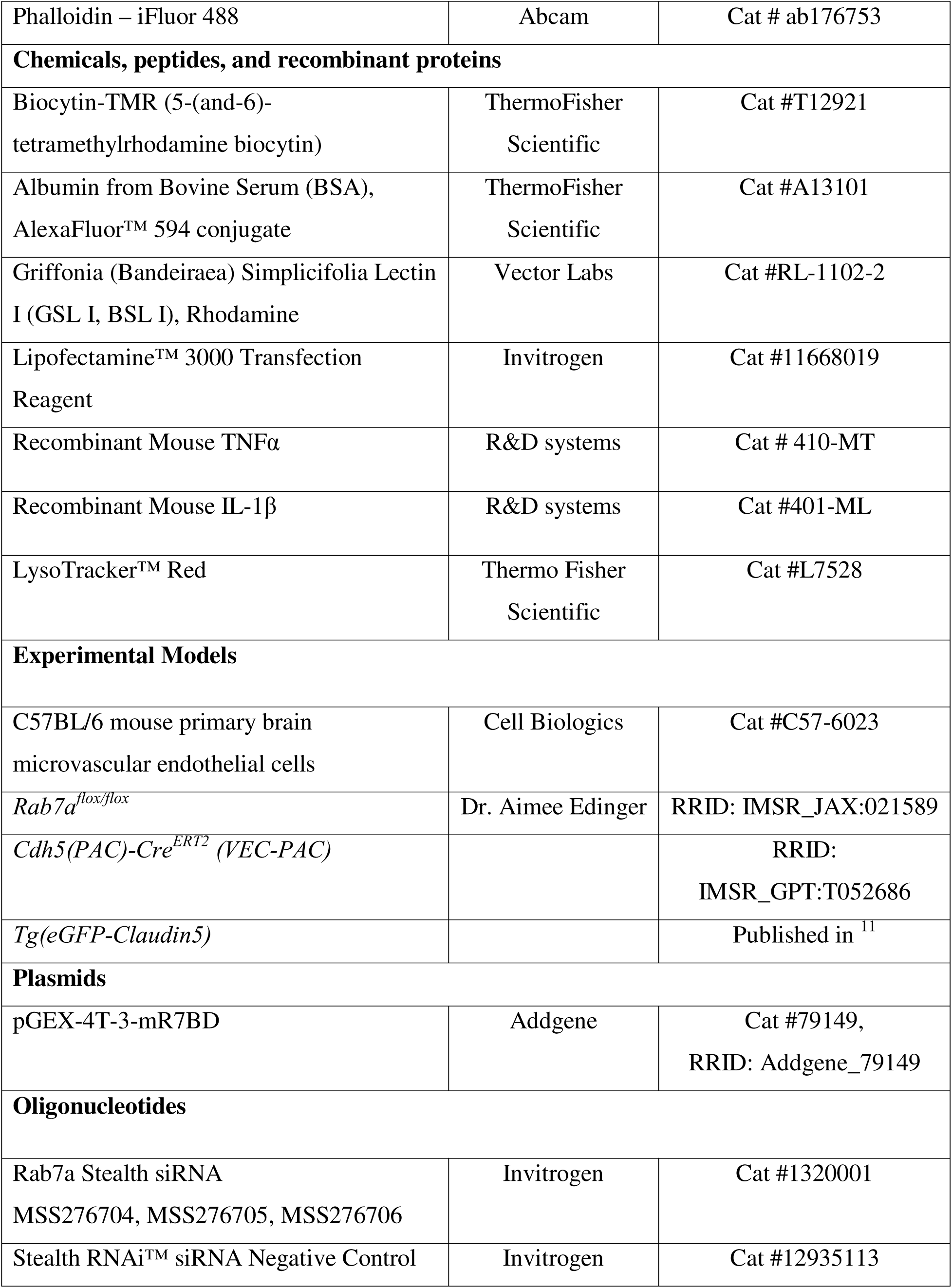
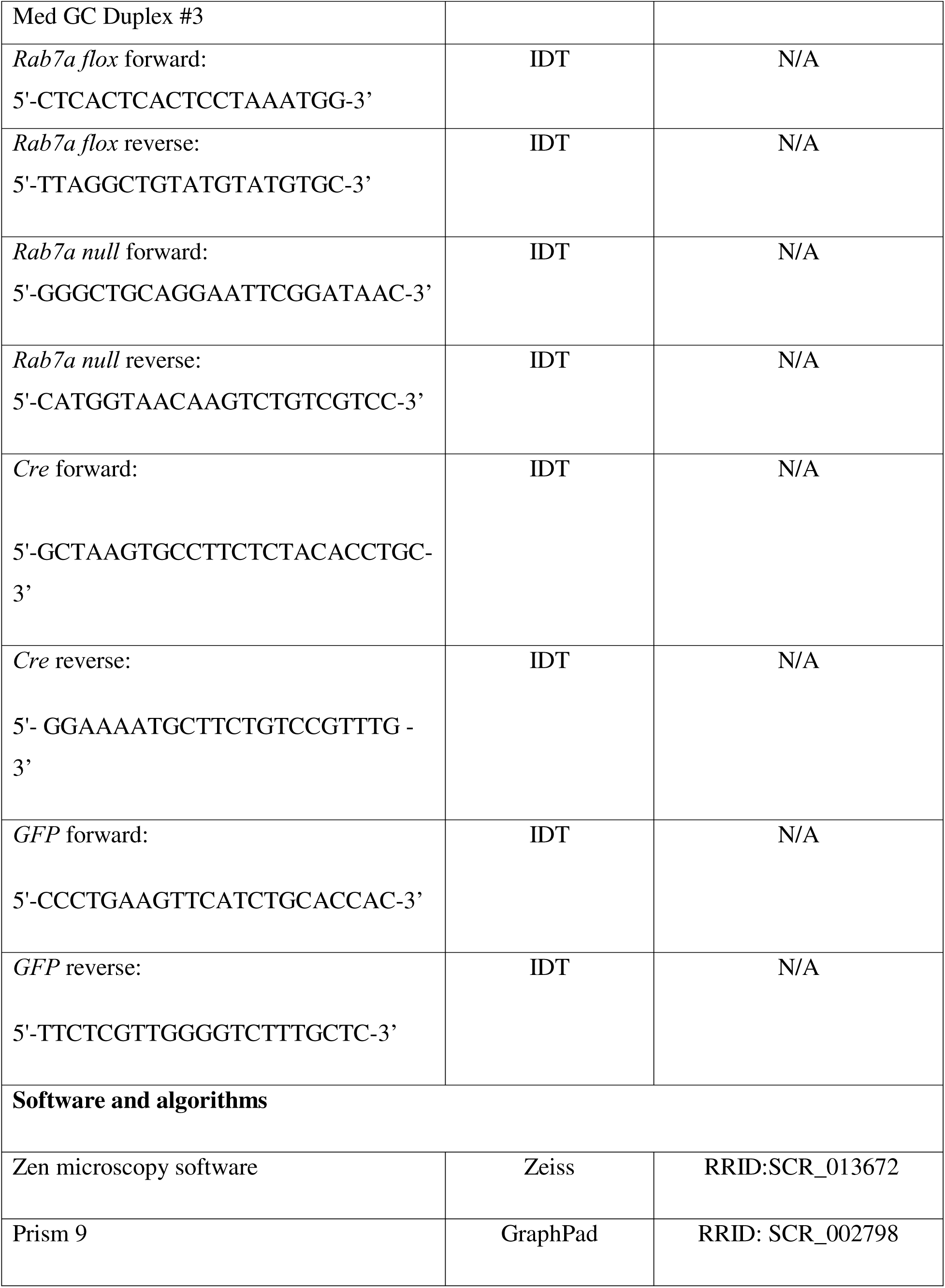
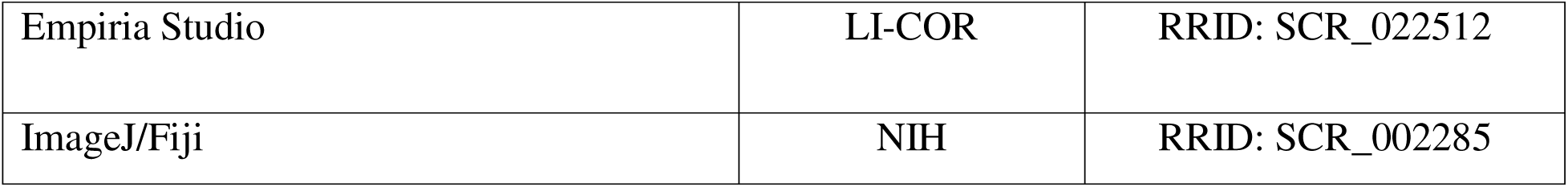

### Methods

#### Mouse ischemic stroke model

Ischemic stroke was induced in 12-14 weeks old male mice by t-MCAO for 45 minutes as described ^62^. Briefly, mice were anesthetized with 1.5-2% isoflurane, left external and common carotid artery were ligated and a 7-0 silicon rubber-coated monofilament (Doccol) was inserted through the common carotid artery into the internal carotid artery. Reperfusion was obtained by removing the monofilament 45 minutes after its insertion. The body temperature of the mice was monitored and maintained at 37°C throughout the procedure. The neurological deficits in mice after t-MCAO were evaluated and scored as described in Jiang S.X. et al., 2005 ^44^.

#### Biocytin-TMR, 70kDa Dextran-TMR and serum IgG permeability analysis

Mice received a tail vein injection of 100 μl of biocytin-TMR (1% in PBS; ThermoFisher Scientific) or 70 kDa dextran-TMR (1% in PBS, ThermofisherScietific). The dye was allowed to circulate for 30-45 minutes. For Biocytin-TMR leakage analysis, mice were anesthetized with isoflurane and perfused first with PBS, then with 4% paraformaldehyde (PFA) in PBS. AlexaFluor 488-conjugated goat anti-mouse IgG (Invitrogen, 1:500) was used to visualize serum IgG leakage in brain sections. The analysis of biocytin-TMR or serum IgG leakage was performed as described before ^11^. Briefly, brain sections were imaged with an upright Zeiss Axioimager fluorescence microscope. Biocytin or IgG leakage was quantified with Fiji software. Brain slices were uniformly thresholded in order to quantify the total area of IgG or biocytin-TMR leakage. Areas that exceeded the threshold levels were defined as leakage area. Average intensity values for biocytin-TMR were gathered by selecting identical regions in ipsilateral or contralateral cortices or livers across subjects, then the ratio between the ipsilateral and contralateral cortex was determined. For 70kDa dextran-TMR analysis, the brains were collected from anesthetized, but not perfused, mice, the ipsilateral and contralateral cortexes were dissected, the tissue was homogenized and centrifugated and the tracer was extracted in PBS as described ^63^. The measurement of the fluorescence from the tissue and quantification were done using a 96-well black plate in an AccuScanFC plate reader (ThermoFisher) with the excitation/emission filters values of 550/570 to detect TMR as described ^63^.

#### Immunofluorescence staining

The brains and livers were dissected from PFA-perfused mice, fixed in 4% PFA at 4°C for 6 hours, washed three times with PBS for 30 minutes per wash, cryoprotected in 30% sucrose/PBS overnight and embedded in Tissue-Tek OCT. Brains were sectioned in 12 μm-thick coronal slices spanning all bregma regions of interest using a Leica cryostat. For immunofluorescence staining of some proteins (e.g. Rab7a) brains were dissected after perfusion of mice with PBS and fresh-frozen in Tissue-Tek. Brain sections were fixed with cold 95% ethanol for 30 minutes and acetone for 1 minute followed by three 5-minute washes with PBS. For immunofluorescence staining of tight junction proteins (Claudin-5, ZO-1) in PFA-fixed brain sections, antigen retrieval was performed with 1X citrate buffer (pH=6, Millipore, Cat # C9999) warmed to 100°C in a water bath for 25 minutes then cooled to room temperature. For primary mBECs cultures, cells were fixed with cold 95% ethanol for 30 minutes and acetone for 1 minute followed by three 5-minute washes with PBS. The following primary antibodies were used for immunofluorescence staining of brain sections or primary mouse BECs: rabbit anti-Rab7a (1:100, Abcam), goat anti-GFAP (1:250, Abcam), rat anti-VE-cadherin (1:25, BD Pharmigen), mouse anti-ZO-1 (1:500, Invitrogen), rabbit anti-ZO-1 (1:500, Invitrogen), rabbit anti-GLUT1 (1:200, Thermo Scientific), rabbit anti-Claudin5 (1:500, Invitrogen), rabbit anti-VE-cadherin (1:250, Abcam), rat anti-CD68 (1:500, Abcam), rabbit anti-Iba1 (1:1000, Wako), mouse anti-NeuN (1:500, Millipore). BSL-rhodamine (1:250, Vector Laboratories) was used to label the vasculature and Phalloidin-iFluor 488 (1:1000 Abcam) was used to label F-actin. The following were used as secondary antibodies for immunofluorescence staining: goat anti-rat Alexa Fluor 594 (A11007), goat anti-rabbit Alexa Fluor 594 (A11012), goat anti-rat Alexa Fluor 488 (A11006), goat anti-rabbit Alexa Fluor 488 (A11034), donkey anti-rat Alexa Fluor 594 (A21209), donkey anti-rabbit Alexa Fluor 594 (A32754), donkey anti-rat Alexa Fluor 488 (A21208), donkey anti-rabbit Alexa Fluor 488 (A21206), donkey anti-rat Alexa Fluor 647 (A48272), donkey anti-rabbit Alexa Fluor 647 (A32795), donkey anti-goat Alexa Fluor 594 (A32758) and donkey anti-goat Alexa Fluor 488 (A11055) (Thermo Fisher Scientific).

#### Western blotting

Protein levels for endothelial markers were assessed using the Odyssey Sa infrared imaging system (LI-COR). β-actin was used as housekeeping gene. The following primary antibodies were used: mouse anti-Claudin5 (1:500, Invitrogen), rabbit anti-Claudin-5 (1:500, Zymed) rabbit anti-Caveolin-1 (1:2000, Abcam), rabbit anti-Occludin (1:500, Thermofisher), rabbit anti-VE-cadherin (1:500, Abcam), mouse anti-ZO-1 (1:500, Invitrogen), rabbit anti-Rab7a (1:1000, Abcam), mouse anti-GST (1:1000, Santa Cruz), rabbit anti-Ccz1 (1:500, Abcam), rabbit anti-EEA1 (1:2000, Abcam), rabbit anti-LAMP1 (1:1000, Abcam), mouse anti-β-actin (1:10000, Novus Biologicals), p62 (1:10000, Abcam) and Atg5 (1:500, Cell Signaling). IR-Dyes 680 and 800 (1:10000, LI-COR) were used as secondary antibodies. Protein expression levels were quantified using LI-COR Biosciences Image Studio software (version 5.2, 2015) and Empiria Studio software (version 3.2.0.186, 2024), normalized on their respective β-actin expression values and presented as a percentage of protein levels in control conditions.

#### mBEC culture and Rab7a silencing *in vitro*

Primary mBECs were purchased from Cell Biologics and grown in endothelial cell medium supplemented with growth factors and 5% FBS (Cell Biologics). Stealth RNAi duplexes targeting Rab7a were purchased from Invitrogen. Stealth RNAi negative control Med CG Duplex #3 (Invitrogen) was used as negative control in all experiments. 12.5 nM of siRNA was transfected using Lipofectamine 3000 (Invitrogen), following manufacturer’s recommendations.

#### Rab7a-GTP pull-down assay

The plasmid pGEX-4T-3-mR7BD, which expresses a recombinant protein consisting of the Rab7a binding domain of the murine RILP protein fused to the C terminus of GST (Addgene, cat#79149), was transformed into Escherichia coli strain BL21. The transformed bacteria were induced to express the GST-RILP protein, which was then purified from the bacterial cell lysates using glutathione-Sepharose 4B beads (GE Healthcare) as described ^31^. Control and cytokine/OGD treated mBECs were lysed in pull-down buffer (20 mM HEPES, 100 mM NaCl, 5 mM MgCl_2_, 1% TX-100, and protease inhibitors) and sonicated. Protein levels in the mBEC lysates were quantified via the BCA assay. Each pull-down was performed by incubating 30 μl of the GST-RILP bound bead slurry pre-equilibrated in pull-down buffer with 300 μg of cell lysate, and rocking the bead-lysate mixture in a nutator overnight at 4°C. Afterwards, the beads were washed thrice with cold pull-down buffer, and bound proteins were eluted by adding sample loading buffer with SDS and incubating at 95°C for 10 min.

#### Trans-endothelial electrical resistance

mBECs transfected with either siCTRL or siRab7a were plated on poly-D-lysine and collagen IV-coated gold-electrode array 96-well ECIS plates (Applied Biophysics) and grown to confluency. They were then switched to low serum medium (1% FBS) for 24 hours and treated with TNFα and IL1β as described above. The electrical resistance of the cultures was recorded every 20 minutes for the duration of the experiment using an ECIS Z-θ system (Applied Biophysics). GraphPad Prism was used to generate resistance curves and to calculate the area under the curve for each experimental condition, as previously described ^64^.

#### *In vitro* assays with primary mouse BECs

For both albumin uptake and Lysotracker assays, mBECs were grown to confluency on poly-D-lysin- and collagen IV-coated glass-bottom 24-well plates (Greiner Bio-One), switched to low serum (1% FBS) medium and treated with TNFα and IL1β as described above. Cells were incubated with either AlexaFluor 594-albumin (100 μg/ml; Thermo Fisher Scientific) or Lysotracker (50 nM; Thermo Fisher Scientific) for 1 hour at 37° C, washed with PBS and fixed in 4% PFA.

#### Microscope image acquisition and quantification

Images of the whole brain sections were acquired with a Zeiss Axioimager fluorescence microscope, higher resolution images were acquired using a Zeiss LSM700 confocal microscope. For comparison purposes, all images of the same staining were acquired under the same settings. All images were processes using Fiji, and all quantifications were carried out blinded. The areas of Biocytin-TMR or IgG leakage were quantified by uniformly thresholding the brain sections and measuring the area exceeding the threshold in each section. The volume of the leakage was calculated as the product of cross-sectional areas and distance between sections. Intensity of the leakage was determined by sampling identical regions in contralateral and ipsilateral cortex in each animal, normalizing their fluorescence intensity on the average fluorescence intensity of the liver from the same animal and calculating the ratio between the ipsilateral and the contralateral cortex.

To quantify junctional abnormalities, vessel segments with intact TJ strands, TJ strands containing gaps or vessel segments without any TJ strands (absent) were identified as described before ^11^ from 5-6 independent ROIs within a specific region and the percentage of vessel segments with either intact TJ strands, TJ strands with gaps or absent junctional strands over the total number of vessel segments was calculated per image in ImageJ. The vessels coverage of junctional strands per each ROI was calculated by thresholding the junctional marker within the vascular marker after masking in ImageJ. To quantify neuronal viability, we calculated the ratio of NeuN positive cells over the total of cells for each field of view. Albumin uptake and Lysotracker *in vitro* were quantified by measuring the area covered by albumin or lysotracker respectively, and dividing it by the number of cells in the field, as previously described ^64^.

#### Transmission electron microscopy

Mice were perfused with PBS for 5 minutes. The ipsilateral or contralateral cortices were dissected from the perfused brains and placed in fixative solution (4% PFA and 2% glutaraldehyde in 0.1 M sodium cacodylate) overnight at 4°C, followed by washing with buffer (0.1 M sodium cacodylate) and water. Fixed samples were treated with 1% reduced Osmium Tetroxide, dehydrated in an ethanol series, followed by treatment with acetonitrile. Samples were then embedded in LX112 resin (Ladd Research Industries) and sectioned. Sections were contrasted with uranyl acetate and lead citrate for imaging in JEOL JEM-1400 TEM equipped with a Veleta (EMSIS, GmbH) CCD camera in the Microscopy and Image Analysis Core Facility, Weill Cornell Medicine (New York, NY, USA). We acquired 16-22 images per cortex (n=3 mice / genotype) and analyzed 46-55 tight junctions per mouse.

#### Statistical analyses

All statistical analyses were performed using GraphPad Prism version 8.1 or higher. Unless differently specified in the figure legend, data are represented as mean ± s.e.m.. The Shapiro-Wilk test was used to verify the normality of datasets. For dataset showing normal distribution, pairwise comparisons were performed using a two-tailed Student’s t-test and multiple comparisons were performed using ordinary one-way ANOVA with Tukey’s multiple comparison test. P values lower than 0.05 were considered statistically significant (***: p<0.001; **: p<0.01; *: p<0.05).

## ACKNOWLEDGEMENTS

We thank Martin Hsu (UNC) and Julian Smith (University of Washington) for initial help with the generation of *Rab7a^iECKO^* mice and the start of the project, Jason Neal for advising with the Rab7a pulldown experiments on primary mBEC lysates and Tyler Cutforth for scientific feedback on this project. A.C., D.J., S.S., M.G., G.P. and D.A. have been supported by grants from the National Eye Institute (R01EY033994), National Heart Lung and Blood Institute (R61/R33 HL159949), National Institute on Aging (RF1AG078352), National Institute of Neurological Disorders and Stroke (R21NS130265), the Leducq Foundation (FDNLEDQ 15CVD 02) and in part by a gift donation from the Ju Foundation. Development of the *Rab7a^fl/fl^* mice was supported by a grant from the National Cancer Institute to ALE (K08 CA100526).

## AUTHOR CONTRIBUTIONS

Conceptualization: A.C., D.A.; Methodology: A.C., D.J., A.A., S.S., M.C.T., M.G., G.P., A. L. E., D.A.; Formal analysis: A.C., D.J., S.S., M.C.T.; Investigation: A.C., D.J., S.S., M.C.T.; Resources: A.L.E, D.A.; Data curation: A.C., D.J., S.S., M.C.T.; Writing - original draft: A.C., D.A.; Writing - revisions: D.J. D.A., Writing feedback: S.S., A.A., M.C.T., A.L.E; Visualization: A.C. D.J.; Supervision: D.A. A.A., Project administration: D.A., Funding acquisition: D.A.

## DECLARATION OF INTEREST

The authors declare no competing or financial interests relevant to the subject matter in this contribution.

## DATA AVAILABILITY STATEMENT

All data are available in the manuscript and the supplementary information. The codes used in this study have been previously published and are freely available (see Key Resources Table). Any additional information required to reanalyze the data reported in this study will be available upon request.

## Notes

### Competing Interest Statement

The authors have declared no competing interest.

### Summary of Updates

The revisions include the following: a) a detailed analysis of structural abnormalities in the subcellular localization of adherens (VE-Cadherin) and tight junction (Claudin-5 and ZO-1)-associated proteins in the blood-brain barrier (BBB) at 48 hours after t-MCAO by immunofluorescence analysis. b) addition of new mice from both Rab7afl/fl and Rab7aiECKO strains to assess changes in total adherens and tight junction protein levels at 48 hours after t-MCAO in the ipsilateral versus contralateral stroke regions using LICOR Western Blotting and quantification. c) Transmission electron microscopy to analyze the structural morphology of tight junctions in the ipsilateral stroke cortical region in Rab7afl/fl and Rab7aiECKO strains at 48 hours after t-MCAO. d) Assessment of BBB permeability to 70-kDa dextran TMR at 48 hours after t-MCAO in both mouse strains as requested by the reviewers. e) Analysis of glial cells [myeloid cell (microglia and macrophages) activation and astrocyte reactivity] at 48 hours after t-MCAO in both mouse strains. f) Additional analyses to assess the effects of Rab7a knockdown on adherens and tight junction proteins in mouse primary brain endothelial cells (BECs) in the absence or presence of pro-inflammatory cytokines (IL-1b and TNFa). g) Addition of an mCLING-Atto 488 assay to measure the internalization rate of VE-Cadherin in siCTRL- and siRab7-transfected mBECs.

