## Supplementary material for "Rab7a is required to degrade select blood-brain barrier junctional proteins after ischemic stroke": Suplementary Information

### SUPPLEMENTARY INFORMATION

#### SUPPLEMENTARY FIGURES AND FIGURE LEGENDS

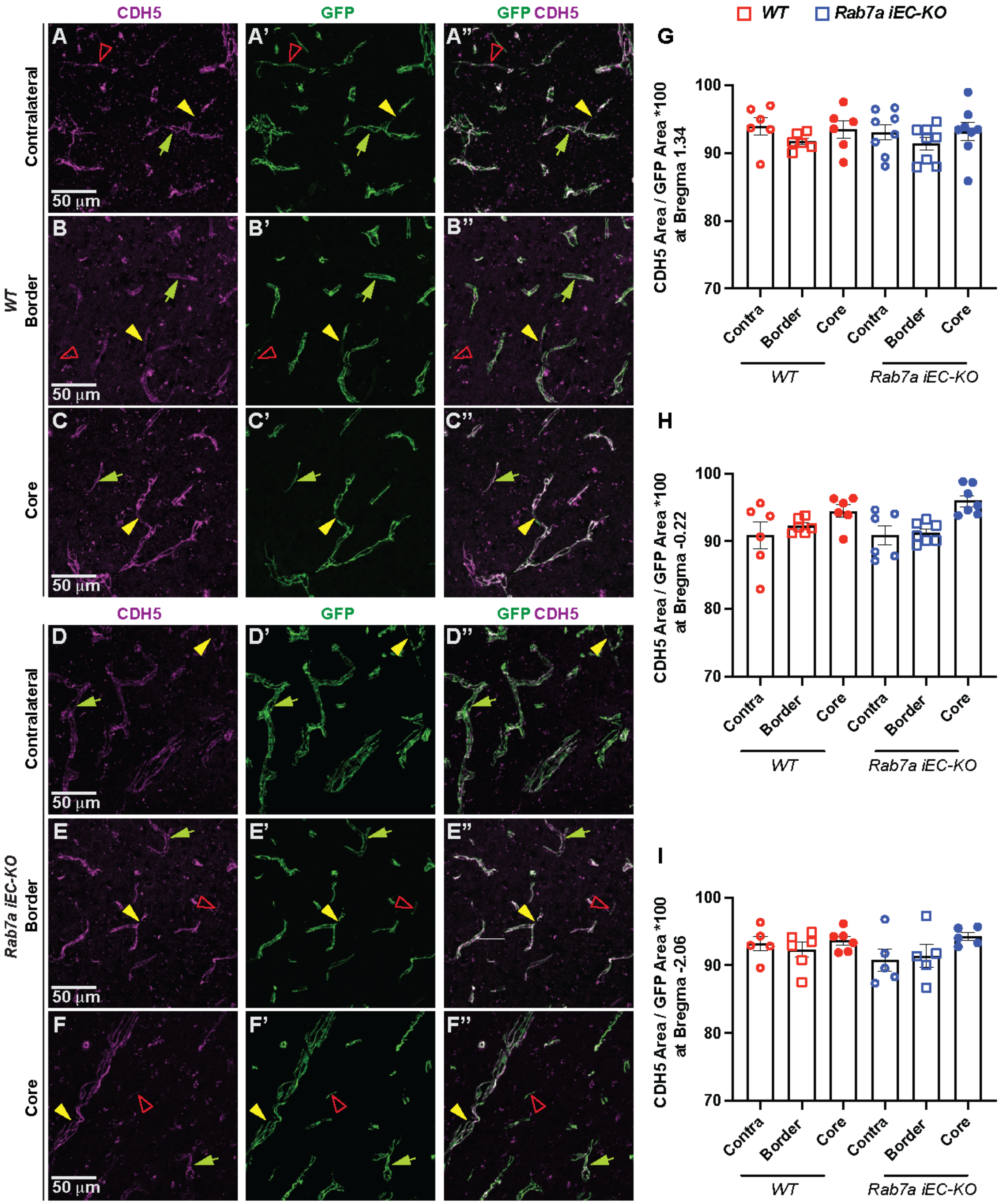

**Figure S1. VE-Cadherin exhibits a similar behavior to eGFP::Claudin-5 in the ipsilateral cortical vessels of *Rab7a*$^{iECKO}$ mice at 48 hours after ischemic stroke.**

**(A-F'')** Immunofluorescences images for VE-Cadherin (CDH5, magenta), and GFP (green) in the contralateral cortex **(A-A'', D-D'')**, border **(B-B'', E-E'')** and core **(C-C'',F-F'')** regions of the ipsilateral cortex of *WT* and *Rab7a*$^{iECKO}$ mice expressing the eGFP::Claudin-5 transgene at 48 hours after t-MCAO. Green arrowheads indicate intact eGFP::Claudin-5$^{+}$ and CDH5$^{+}$ junctional strands, yellow arrowheads point to gaps in eGFP$^{+}$ and CDH5$^{+}$ strands, red open arrowheads indicate absent / intracellular eGFP::Claudin-5$^{+}$ and CDH5$^{+}$ junctional strands. Scale bar: 50 μm. (**G-I**) Quantification of the percentage of the area of CDH5 / area of GFP at bregmas 1.34 **(G)**, -0.22 **(H)**, and -2.06 **(I)** in the contralateral cortex, border and core regions of the ipsilateral cortex of *WT* and *Rab7a*$^{iECKO}$ mice at 48 hours after t-MCAO. Each dot represents an animal (n= 6-8 mice/group). Data are means ± s.e.m. There is no significant difference by one-way ANOVA with post-hoc Tukey's correction.

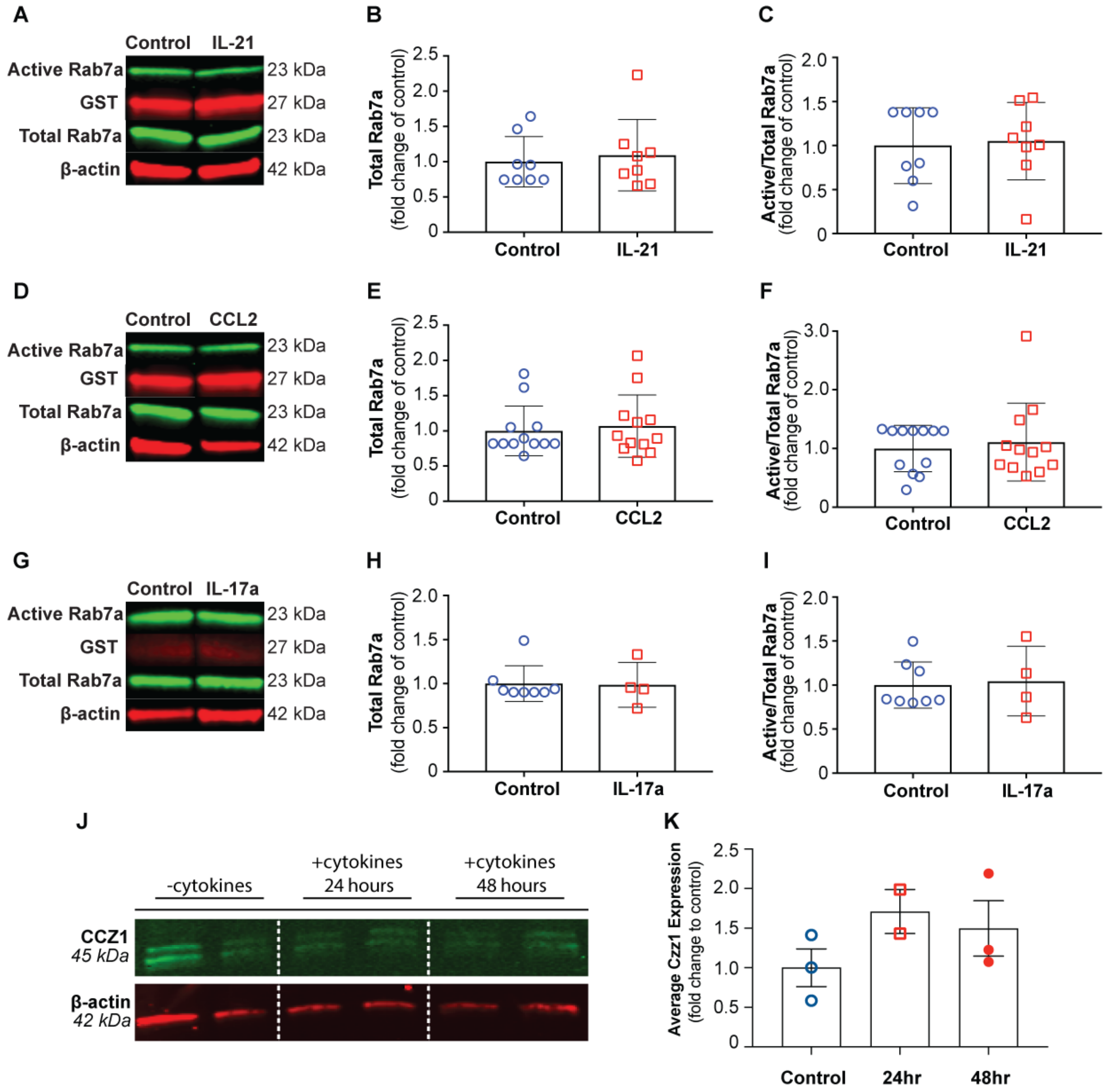


**Figure S2. IL-21, CCL2 and IL-17A do not activate Rab7a in primary mouse brain endothelial cells.**

(**A, D, G**) Western blot of active and total Rab7a protein levels in primary mouse brain endothelial cells (mBECs) treated with IL-21 (10ng/mL) (**A**), CCL2 (10ng/mL) (**D**) and IL-17A (10ng/mL) (**G**) for 48 hours and untreated control cells. GST and β-actin are used to normalize active and total Rab7a levels, respectively. The molecular weight of each protein is shown on the right. (**B, E, H**) Quantification of total Rab7a protein levels in mBECs treated with IL-21 (**B**), CCL2 (**E**), and IL-17A (**H**) for 48 hours and untreated cells. Each dot represents an independent experiment. (**C, F, I**) Quantification of the ratio of active over total Rab7a protein levels in mBECs treated with IL-21 (**C**), CCL2 (**F**) and IL-17A (**I**) for 48 hours versus untreated mBECs. Each dot represents an independent experiment (n.s.: $p>0.05$; Student's t-test). **J, K**) Western blot and quantification of Ccz1 protein levels in primary mBECs treated with IL-1β (10ng/mL) and TNFα (10ng/mL) for 24 and 48 hours and untreated control cells. β-actin is used to normalize protein levels. Ccz1 protein levels are increased, albeit not significantly, between 24 to 48 hours of cytokine treatment. Each dot represents an independent experiment.

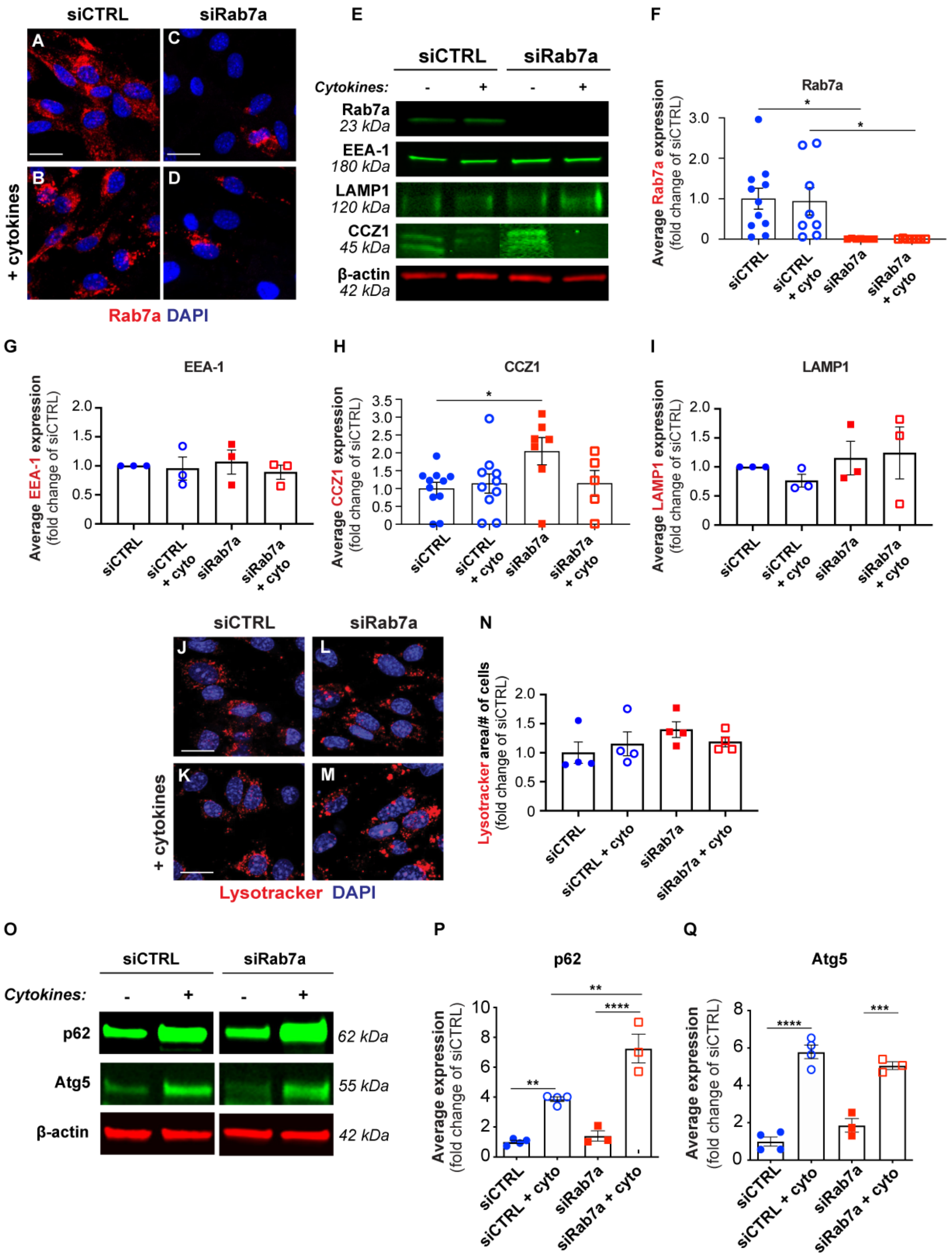
siCTRL
siRab7a
A
C
B
D
+ cytokines
Rab7a DAPI
E
siCTRL
siRab7a
Cytokines:
-
+
-
+
Rab7a
23 kDa
EEA-1
180 kDa
LAMP1
120 kDa
CCZ1
45 kDa
β-actin
42 kDa
F
Rab7a
Average Rab7a expression
(fold change of siCTRL)
3.0
2.0
1.0
0
*
*
siCTRL
siCTRL + cyto
siRab7a
siRab7a + cyto
G
EEA-1
Average EEA-1 expression
(fold change of siCTRL)
2.0
1.5
1.0
0.5
0
siCTRL
siCTRL + cyto
siRab7a
siRab7a + cyto
H
CCZ1
Average CCZ1 expression
(fold change of siCTRL)
3.5
3.0
2.5
2.0
1.5
1.0
0.5
0
*
siCTRL
siCTRL + cyto
siRab7a
siRab7a + cyto
I
LAMP1
Average LAMP1 expression
(fold change of siCTRL)
2.0
1.5
1.0
0.5
0
siCTRL
siCTRL + cyto
siRab7a
siRab7a + cyto
siCTRL
siRab7a
J
L
K
M
+ cytokines
Lysotracker DAPI
N
Lysotracker area/# of cells
(fold change of siCTRL)
2.0
1.5
1.0
0.5
0
siCTRL
siCTRL + cyto
siRab7a
siRab7a + cyto
O
siCTRL
siRab7a
Cytokines:
-
+
-
+
p62
62 kDa
Atg5
55 kDa
β-actin
42 kDa
P
p62
Average expression
(fold change of siCTRL)
10
8
6
4
2
0
**
**
****
siCTRL
siCTRL + cyto
siRab7a
siRab7a + cyto
Q
Atg5
Average expression
(fold change of siCTRL)
8
6
4
2
0
****
***
siCTRL
siCTRL + cyto
siRab7a
siRab7a + cyto

**Figure S3. Rab7a knockdown in primary mouse brain endothelial cells does not affect other proteins of the endolysosomal pathway.**

(**A-D**) Immunofluorescence for Rab7a protein (red) and DAPI (blue) in mBECs transfected with either siCTRL (**A, B**), or siRab7a (**C, D**) without/with pro-inflammatory cytokines (IL-1$\beta$ =10ng/mL and TNF-$\alpha$ = 10ng/mL). Rab7a protein levels are almost negligent after siRab7a transfection. Scale bar: 10 µm. **(E-I**) Western blots **(E)** and quantification for Rab7a **(F)**, EEA-1 (**G**), Ccz1 **(H)** and LAMP1 **(I)** proteins in siCTRL- and siRab7a-transfected mBECs without/with cytokines (each dot represents an independent experiment; data are means ± s.e.m.; *: $p<0.05$; n.s.: $p>0.05$; one-way ANOVA with post-hoc Tukey's correction). (**I-L**) Representative fluorescence images and (**M**) quantification of lysotracker (red) in siCTRL- (**I, J**) and siRab7a- (**K, L**) transfected mBECs without/with cytokine treatment. DAPI (blue) labels nuclei. Rab7a knockdown has no effect on the number of lysosomes per mBEC. Scale bar: 10 µm. Data are means ± s.e.m.; n.s. $p>0.05$ (not shown); one-way ANOVA with post-hoc Tukey's correction. (**O-Q**) Western blots and quantification for p62, Atg5 and $\beta$-actin (control) proteins in siCTRL- and siRab7a-transfected mBECs without/with pro-inflammatory cytokines. Each dot represents an independent experiment. Cytokine treatment increases autophagy protein levels, and Rab7a knockdown increases p62 protein accumulation after cytokine treatment indicative of reduced autophagosome degradation. Each dot represents an independent experiment. Data are means ± s.e.m.; **** $p<0.0001$; *** $p<0.001$ ** $p<0.01$; *: $p<0.05$; n.s. $p>0.05$ (not shown); one-way ANOVA with post-hoc Tukey's correction.

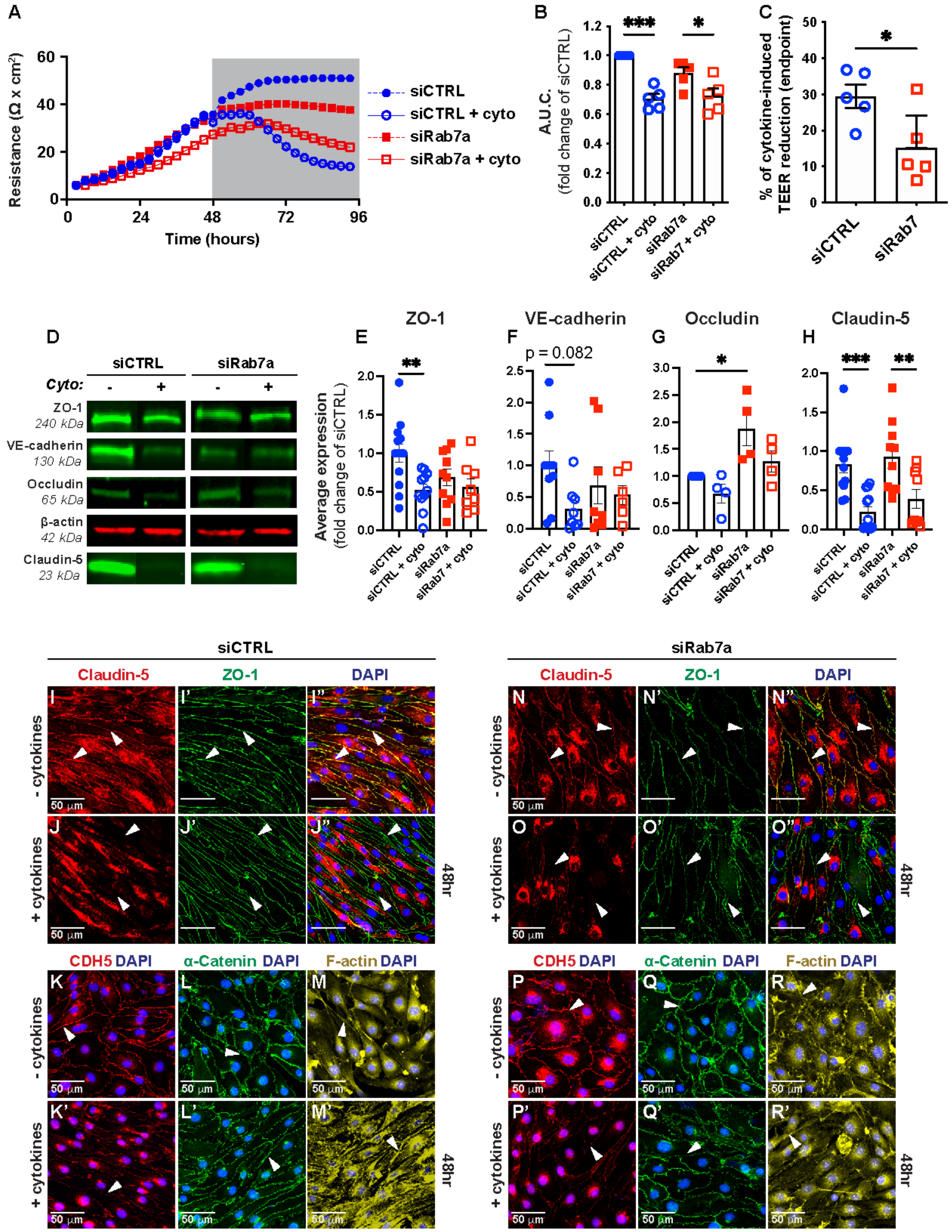
A
Resistance (Ω x cm²)
Time (hours)
siCTRL
siCTRL + cyto
siRab7a
siRab7a + cyto
B
A.U.C.
(fold change of siCTRL)
***
*
siCTRL
siCTRL + cyto
siRab7a
siRab7 + cyto
C
% of cytokine-induced TEER reduction (endpoint)
*
siCTRL
siRab7
D
siCTRL
siRab7a
Cyto:
ZO-1 240 kDa
VE-cadherin 130 kDa
Occludin 65 kDa
β-actin 42 kDa
Claudin-5 23 kDa
E
ZO-1
**
Average expression
(fold change of siCTRL)
F
VE-cadherin
p = 0.082
G
Occludin
*
H
Claudin-5
***
**
siCTRL
siCTRL + cyto
siRab7a
siRab7 + cyto
siCTRL
siRab7a
Claudin-5
ZO-1
DAPI
- cytokines
+ cytokines
48hr
50 µm
CDH5 DAPI
α-Catenin DAPI
F-actin DAPI

**Figure S4. Rab7a knockdown reduces cytokine-driven BEC barrier dysfunction by reducing degradation of select junctional proteins and affects reorganization of F-actin bundles at cell junctions under inflammatory conditions in mouse brain endothelial cells.**

**(A)** Transendothelial electrical resistance (TEER) measurement with the ECIS instrument of primary mouse brain endothelial cells (mBECs) transfected with either an siRNA targeting Rab7a (siRab7a), or a scrambled siRNA (siCTRL) without/with inflammatory cytokines (TNFα, IL-1β 10 ng/mL). The gray area represents the period of time during which cells were treated with cytokines or vehicle. (**B, C**) Quantification of the area under the curve (A.U.C.) during treatment with cytokines [gray area in (a)] and of the percentage of TEER reduction 48 hours after treatment with cytokines (right panel). Each dot represents an independent experiment. Data are means ± s.e.m. **: $p<0.01$; *: $p<0.05$; n.s. $p>0.05$ (not shown); one-way ANOVA with post-hoc Tukey's correction (left graph), and Student's t-test (right graph). **(D)** Western blot for junctional proteins ZO-1, VE-Cadherin, Occludin, and Claudin-5 in siCTRL- and siRab7a-transfected mBECs without/with cytokines. β-actin serves as control. The respective molecular weights of the proteins are shown under the name. **(E-H)** Quantification of ZO-1, VE-Cadherin, Occludin, and Claudin-5 protein levels in siCTRL- and siRab7a-transfected without/with cytokines. Each dot represents an independent experiment from 3 biological replicates. Data are means ± s.e.m.; ***: $p<0.001$; **: $p<0.01$; *: $p<0.05$; one-way ANOVA with post-hoc Tukey's correction. (**I-R'**) Immunofluorescence staining of mBECs transfected with either siCTRL (**I-M'**) or siRab7a (**N-R'**) without/with cytokine treatment for the following proteins: Claudin-5 (red), ZO-1 (green), and DAPI (nuclei, blue) [siCTRL (**I-J")** and siRab7a (**N-O"**)], CDH5 (red) and DAPI (blue) (**K, K'** and **P, P'**), α-Catenin (green) and DAPI (blue) (**L, L'** and **Q, Q'**), F-actin (yellow) and DAPI (blue) (**M, M'** and **R, R'**). White arrowhead indicates the presence of the protein at cell junctions in mBECs. Scale bar: 50 µm.

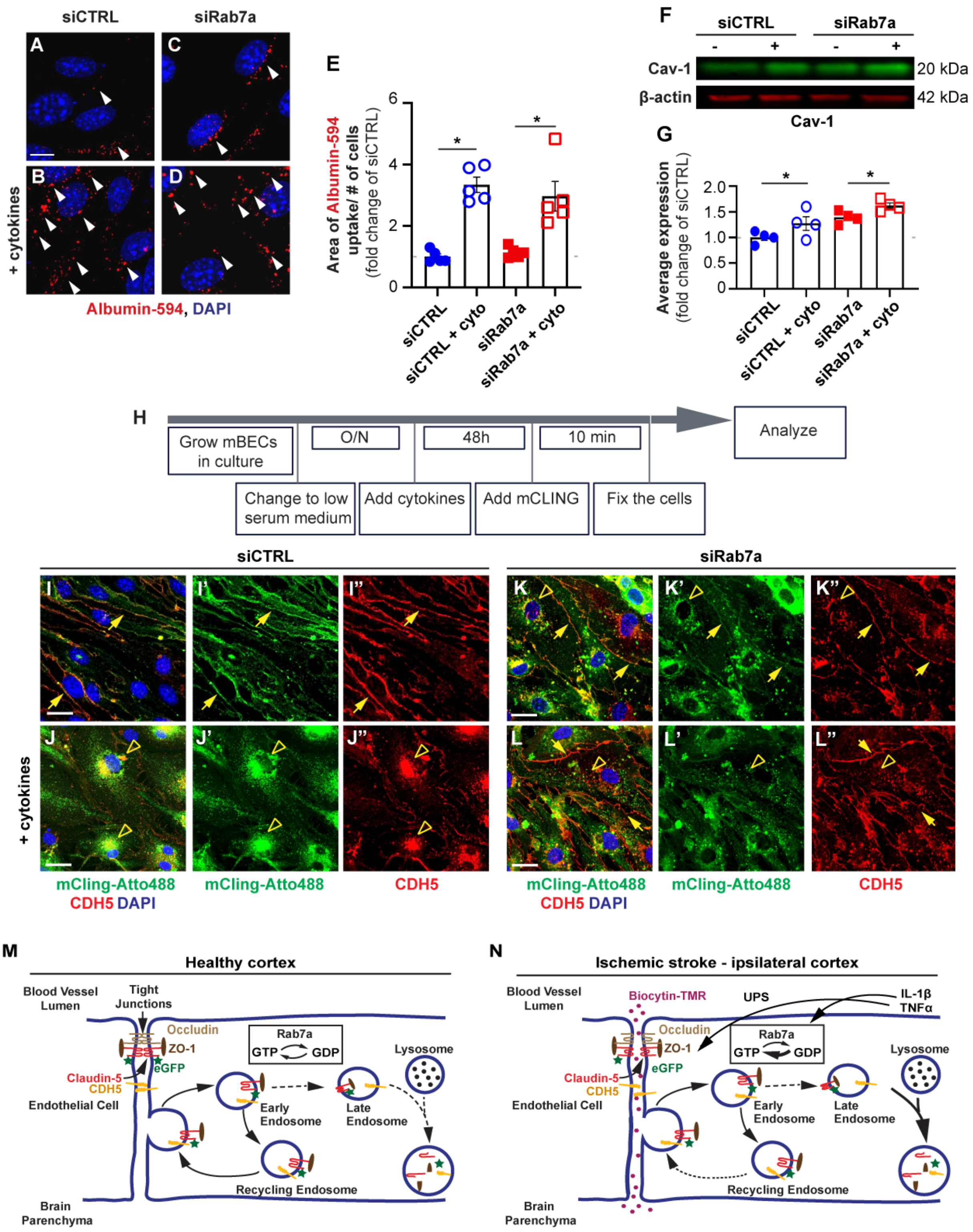
siCTRL
siRab7a
A
B
C
D
+ cytokines
Albumin-594, DAPI
E
Area of Albumin-594 uptake/ # of cells
(fold change of siCTRL)
0
2
4
6
*
*
siCTRL
siCTRL + cyto
siRab7a
siRab7a + cyto
F
siCTRL
siRab7a
-
+
-
+
Cav-1
β-actin
20 kDa
42 kDa
Cav-1
G
Average expression
(fold change of siCTRL)
0
0.5
1.0
1.5
2.0
*
*
siCTRL
siCTRL + cyto
siRab7a
siRab7a + cyto
H
Grow mBECs in culture
O/N
48h
10 min
Analyze
Change to low serum medium
Add cytokines
Add mCLING
Fix the cells
siCTRL
siRab7a
I
I'
I''
J
J'
J''
K
K'
K''
L
L'
L''
+ cytokines
mCling-Atto488 CDH5 DAPI
mCling-Atto488
CDH5
mCling-Atto488 CDH5 DAPI
mCling-Atto488
CDH5
M
Healthy cortex
Blood Vessel Lumen
Tight Junctions
Occludin
ZO-1
eGFP
Claudin-5
CDH5
Endothelial Cell
Rab7a
GTP
GDP
Lysosome
Early Endosome
Late Endosome
Recycling Endosome
Brain Parenchyma
N
Ischemic stroke - ipsilateral cortex
Blood Vessel Lumen
Biocytin-TMR
UPS
IL-1β
TNFα
Occludin
ZO-1
eGFP
Claudin-5
CDH5
Endothelial Cell
Rab7a
GTP
GDP
Lysosome
Early Endosome
Late Endosome
Recycling Endosome
Brain Parenchyma

**Figure S5. Rab7a knockdown in primary mouse brain endothelial cells does not affect caveolar-mediated uptake of albumin, but increases CDH5 internalization rate after cytokine treatment.**

**(A-D)** Immunofluorescence for the uptake of albumin-Alexa594 (red; white arrowheads) in siCTRL or siRab7a-transfected mBECs with/without cytokines. DAPI (blue) labels nuclei. **(E)** Quantification of albumin-Alexa 594 uptake by siCTRL- or siRab7a-transfected mBECs without/with cytokines. Each dot represents an independent experiment. Data are means $\pm$ s.e.m.; *: $p<0.05$; n.s. $p>0.05$ (not shown); one-way ANOVA with post-hoc Tukey's correction. Rab7 knockdown has no effect on cytokine-induced uptake of albumin-Alexa594. **(F, G)** Western blot analysis and quantification of Caveolin-1 expression in siCTRL- or siRab7a-transfected mBECs with or without cytokines (Each dot represents an independent experiment; data are means $\pm$ s.e.m.; *: $p<0.05$; n.s. $p>0.05$ (not shown); one-way ANOVA with post-hoc Tukey's correction). **(H)** Diagram of experimental design for the mCLING-Atto488 assay. **(I-L'')** Immunofluorescence for mCling-Atto488 (green), CDH5 (red), and DAPI (blue) in mBECs transfected with either siCTRL **(I-J'')** or siRab7a (**K-L''**) after 48 hours without/with cytokines. Normal arrowhead indicates labelled protein present at junctions and absent arrowheads indicates internalized protein. **(M-N)** Working model for the role of Rab7a in regulation of select BBB junctional proteins after ischemic stroke. In healthy conditions, Rab7a cycles equally between its active and inactive states, balancing the rates of degradation and recycling of internalized adherens (VE-Cadherin, CDH5, yellow) and tight junction [Claudin-5 (red) and ZO-1 (brown)] associated proteins. Under pathological conditions, such as ischemic stroke, some inflammatory cytokines (e.g. IL-1$\beta$, TNF$\alpha$) activate Rab7a, although the mechanism remains unclear. The activated Rab7a increases the rate of degradation of select junctional proteins in BECs leading to increased paracellular BBB permeability after ischemic stroke. Rab7a-independent mechanisms also contribute to degradation of BBB junctional proteins at the acute phase of ischemic stroke.
